# Proteome-wide C-degron activity profiling connects conditional regulation of the CTLH E3 ligase complex to ribosome biogenesis

**DOI:** 10.64898/2026.01.14.698769

**Authors:** Drew W. Grant, Shengjiang Tan, Dylan E. Ramage, Mamie Z. Li, Ying Di, Iva A. Tchasovnikarova, Michael P. Weekes, Stephen J. Elledge, Alan J. Warren, Richard T. Timms

## Abstract

Many E3 ubiquitin ligases recognize cognate degron motifs located at protein termini, but the paucity of *bona fide* substrates of N-degron and C-degron pathways hampers our understanding of their physiological significance. Here, by devising an expression screening approach to assess the effect of C-terminal “capping” on the stability of thousands of human proteins, we systematically identify a suite of full-length substrates harboring C-terminal degrons. Interrogating one leading candidate, ZMYND19, we characterize a C-degron pathway governed by the Muskelin substrate adaptor of the CTLH E3 ligase complex. Cell-to-cell variability in ZMYND19 stability uncovered conditional regulation, with CTLH-mediated degradation impaired by TNF-α stimulation but enhanced by mTOR inhibition. Parallel genetic and proteomic screens identified two poorly characterized proteins, AAMP and AEN, as additional substrates of the CTLH^Muskelin^ C-degron pathway, leading us to define an essential role for AAMP in ribosome maturation through chaperone activity towards ribosomal protein uL16. Altogether, these data define a C-degron pathway through which the Muskelin substrate adaptor connects conditional regulation of the CTLH E3 ligase complex to control of ribosome biogenesis.

## INTRODUCTION

The ubiquitin-proteasome system (UPS) represents the primary route through which eukaryotic cells achieve selective protein degradation^1^. As such, the UPS plays a critical role in the regulation of all major cellular processes, with tasks ranging from quality control of misfolded proteins to maintenance protein homeostasis through to dynamic protein turnover in response to changing environmental conditions^2,3^. Ubiquitin is a 76-amino acid protein, which, through the sequential action of E1, E2 and E3 enzymes, is covalently conjugated to substrate proteins^4,5^. The potential biological consequences of protein ubiquitination are diverse^6^, but the formation of polyubiquitin chains formed via lysine-48 (K48) linkages canonically serve as a targeting signal to the 26S proteasome for proteolytic degradation^7,8^. The importance of this process is underscored by the dysregulation of the UPS that is observed in a variety of disease states, including cancer and neurodegeneration^9,10^.

A major unresolved question concerns how the UPS achieves such high selectivity towards its myriad of substrates. The primary specificity determinants are the E3 ubiquitin ligases, which are typically responsible for the recruitment of both the target substrate and an E2 ubiquitin conjugating enzyme to catalyze ubiquitination. Substrate identification involves an E3 ligase binding to a cognate degron, defined as the minimal element within a substrate protein sufficient to allow E3 ligase recognition^11^. Known degrons exhibit considerable diversity: they can comprise short peptide motifs lying within unstructured regions of proteins, or extended interfaces with elaborate three-dimensional structures; they can also act either constitutively or, such as those created (or destroyed) by post-translational modifications, in a conditional manner^12^. Currently, however, it is only for a very small proportion of the >600 E3 ubiquitin ligases encoded in the human genome that we have a molecular understanding of the degron motifs that they target^13,14^, and overcoming this bottleneck represents a key challenge towards the goal of achieving a systems-level understanding of UPS function.

In principle, degrons may lie anywhere in the substrate protein. Protein termini appear particularly enriched for degrons, however, with a growing number of E3 ligases now defined to recognize short sequence motifs specifically when located at or near the ends of proteins^15,16^. Indeed, the first degrons to be discovered in the 1980s lay at the extreme N-terminus of proteins^17^, a discovery which gave rise to the study of the N-degron (formerly N-end rule^18^) pathways over the past forty years. However, it was much more recently that an analogous set of C-degron pathways were first identified^19,20^. These include the Kelch family of Cul2 substrate adaptors which recognize C-terminal glycine degrons^21^, the FEM family of Cul2 substrate adaptors which recognize C-terminal arginine or proline degrons^22,23^, and TRIM7 which targets C-terminal glutamine^24–26^. However, although we are beginning to delineate degron motifs at protein C-termini that confer instability, in most cases we still have little understanding of the key physiological substrates of these pathways and hence the cellular context in which they operate. Although some may serve roles in quality control, for example Cul2^KLHDC10^ and Pirh2 recognize alanine tails in ribosome quality control^27^ and Cul4^CRBN^ recognizes C-terminal imides that can arise from intramolecular cyclisation of asparagine and glutamine^28^, it is likely that others regulate specific cellular processes. Furthermore, although we and others have successfully defined a broad spectrum of C-terminal degron motifs on the basis of their effects in the context of short peptides fused to reporter proteins such as GFP^20,29,30^, this approach may not necessarily identify E3 ligase-substrate associations that are of physiological significance in the context of the corresponding endogenous proteins.

To overcome this limitation, here we devised an expression screening approach to measure the impact of manipulating C-terminal sequences on the stability of full-length proteins. Through comparative stability profiling of two human ORFeome libraries, we systematically assessed the effect of C-terminal “capping” on the stability of thousands of human proteins. Of the ∼50 high-confidence substrates bearing putative C-degrons, our follow-up work focused on one of the leading candidates, ZMYND19, leading us to define a C-degron pathway through which the kelch repeats of the Muskelin substrate adaptor of the CTLH E3 ligases complex recognizes a -PER* C-terminal degron. We show that this C-degron pathway is conditionally regulated, with TNF-α stimulation inhibiting ZMYND19 degradation and mTOR inhibition enhancing it, and through parallel genetic and proteomic approaches identify AAMP and AEN as additional substrates. Finally, we define the hitherto uncharacterized protein AAMP as the ortholog of the yeast ribosome assembly factor Sqt1, showing that it is required for the function of the large ribosomal subunit protein uL16 and thus connecting CTLH to the control of ribosome biogenesis. Altogether these data delineate a novel C-degron pathway regulated by the CTLH complex, characterize the physiological circumstances in which the pathway acts, and provide a resource permitting the characterization of additional C-degron pathways in the future.

## RESULTS

### Systematic identification of full-length proteins harboring C-terminal degrons

With the goal of identifying novel C-degron pathways which regulate the stability of full-length proteins, we utilized the dual-color Global Protein Stability (GPS) expression screening platform^31^ to measure protein stability proteome-wide. Two distinct human ORFeome libraries were fused downstream of GFP. Previously we described the use of the Ultimate ORFeome library (hereafter “uORFeome”) for stability profiling^20^, which comprises ∼17,000 human open reading frames (ORFs) which retain stop codons at their native positions **(Fig. 1A)**. Here we generated an analogous platform using the human ORFeome V8.1 library, which encodes ∼13,000 human ORFs which lack stop codons; in the context of the GPS-ORFeome V8.1 expression vector, this results in proteins which terminate with an invariant 8-mer peptide (-PTFLYKVV*) encoded by the vector backbone **(Fig. 1B)**. We reasoned that, for the ∼8,000 proteins in common between the two libraries **(Fig. S1A)**, comparison of stability profiles in the uORFeome library (native end) versus the ORFeome V8.1 library (“capped” end) would identify proteins whose stability was determined via C-terminal motifs. We hypothesized that “capping” of the C-terminus would obscure recognition motifs for E3 ligases targeting C-terminal degrons, such that substrates subject to constitutive degradation via C-degron pathways would exhibit increased stability in the context of the ORFeome V8.1 library **(Fig. S1B)**.

**Figure 1.**
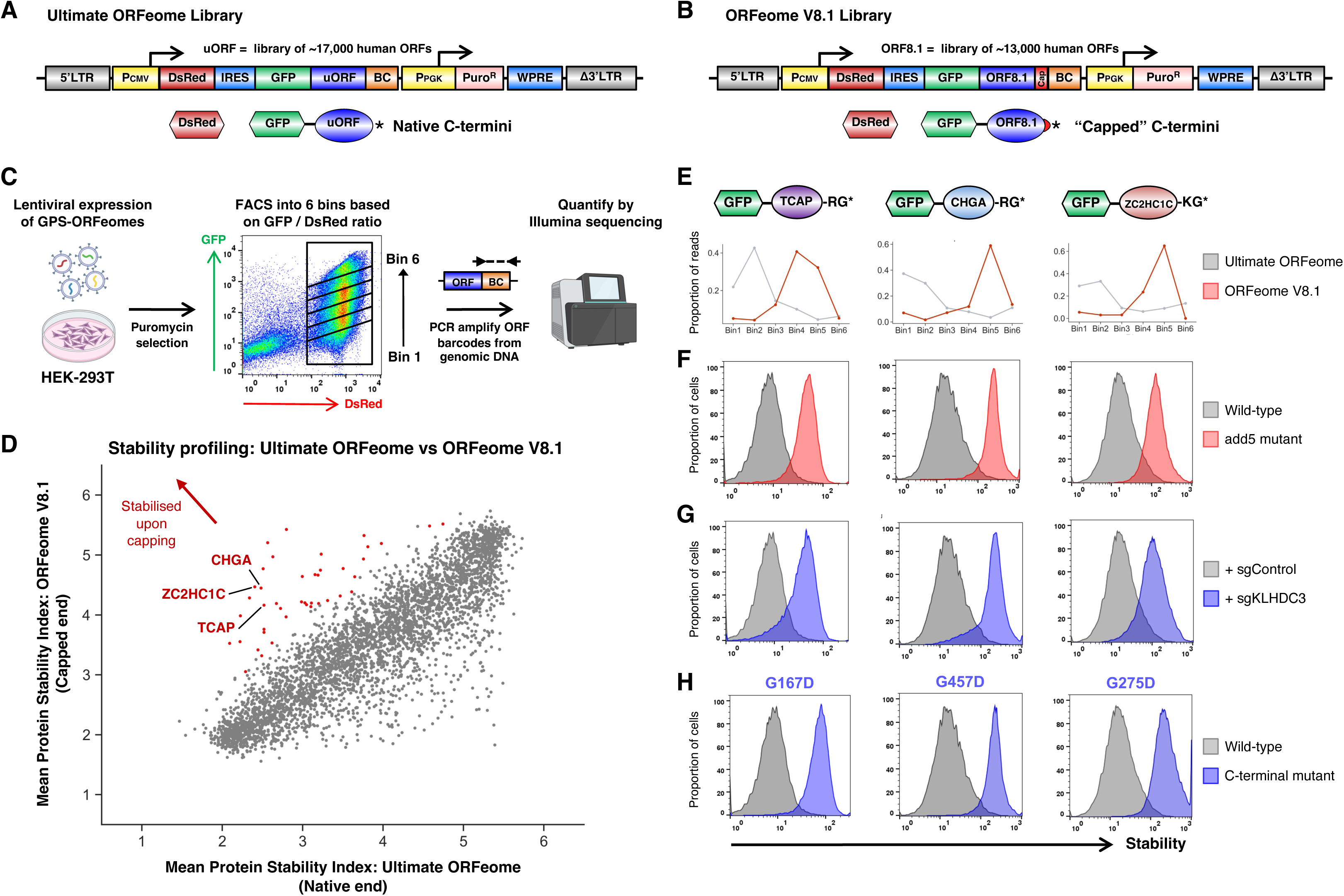
Comparative stability profiling identifies full-length human proteins bearing C-terminal degrons. **(A-B)** Schematic representation of the two human ORFeome libraries. The Ultimate ORFeome library **(A)** encodes proteins bearing stop codons at their native position, whereas the ORFeome V8.1 **(B)** encodes proteins ‘capped’ with an invariant 9 additional amino acids at their C-terminus. (BC, barcode; WPRE, Woodchuck Hepatitis Virus Posttranscriptional Regulatory Element; LTR, long terminal repeat) **(C)** Schematic representation of the comparative stability profiling screen. The two ORFeome libraries were mixed, packaged into lentiviral particles and introduced at single copy into HEK-293T cells; stability profiling was performed by partitioning the population into six stability bins of equal size by FACS, followed by PCR amplification of the ORF barcodes from genomic DNA and quantitation by Illumina sequencing. **(D)** Comparative stability profiling identifies proteins harboring C-terminal stability determinants. Each dot represents the mean stability score (1 = maximally unstable, 6 = maximally stable) for each protein in common between the two ORFeome libraries, with red dots indicating proteins exhibiting stabilization >0.75 PSI units in the ORFeome V8.1 library across both replicate experiments. **(E-H)** Successful identification of positive control substrates harboring glycine C-degrons targeted by Cul2^KLHDC3^. **(E)** Screen profiles for TCAP, CHGA and ZC2HC1C (isoform 2), reflecting the distribution of sequencing reads across the six stability bins when assayed as part of the Ultimate ORFeome library (gray) or the ORFeome V8.1 library (red). **(F)** “Capping” of the C-terminus of each protein with a five amino acid peptide (-KASTN*; “add5”) resulted in stabilization. Each protein also exhibited stabilization upon CRISPR/Cas9-mediated targeting of *KLHDC3* **(G)** or mutation of the glycine residue at the extreme C-terminus **(H)**.

We combined the two GPS-ORFeome libraries, packaged them into lentiviral particles, and transduced HEK-293T cells at low multiplicity of infection (∼0.2) such that the overwhelming majority of cells expressed at most one single GFP-fusion construct. Following puromycin selection to eliminate untransduced cells, we performed stability profiling by FACS coupled with Illumina sequencing **(Fig. 1C)**. Reassuringly, most proteins exhibited equivalent stability irrespective of their library of origin, with the ∼5,000 proteins quantified in both libraries showing highly concordant Protein Stability Index (PSI) scores **(Fig. 1D and Table S1)**. Moreover, this effect was highly reproducible across duplicate experiments **(Fig. S1C)**. However, a considerable number of proteins did exhibit differential stability across the two libraries. Among the 52 proteins that were concordantly stabilized >0.75 PSI units in the ORFeome V8.1 library compared to the uORFeome library across both replicate experiments, we found several examples of proteins previously known to be targeted for proteasomal degradation via C-degron motifs when expressed in the context of the GPS system **(Fig. 1D and Fig. S1C)**. These included titin-cap (TCAP), chromogramin-A (CHGA), and isoform 2 of ZC2HC1C (formerly known as FAM164C), which all terminate with C-terminal glycine degrons^20^ **(Fig. 1E and Fig. S1D)**. In each case we verified that the stabilization we observed in the context of the ORFeome V8.1 library was not specific to the sequence of the 8-mer peptide appended, as addition of a “neutral” 5-mer peptide^20^ (-KASTN*; “add5”) to the C-terminus also resulted in stabilization **(Fig. 1F)**. We also confirmed that the activity of Cul2^KLHDC3^ was indeed responsible for their instability, as CRISPR/Cas9-mediated disruption of *KLHDC3* **(Fig. 1G)** or substitution of the C-terminal glycine residue **(Fig. 1H)** also resulted in stabilization. Altogether, we concluded that our comparative GPS-ORFeome screens successfully identified full-length proteins subject to degradation via C-degron motifs, defining a set of ∼50 proteins harboring putative C-degrons for detailed characterization **(Fig. S2)**.

### ZMYND19 harbors a C-terminal degron

We focused our follow-up studies on one of the leading candidates, ZMYND19, which exhibited markedly greater stability upon C-terminal capping in the ORFeome V8.1 library compared to the uORFeome library **(Fig. 2A and Fig. S1D)**. The function of ZMYND19 remains unknown, with its only annotated feature a MYND-type zinc finger domain. We first confirmed that its stability was indeed regulated via its C-terminus, as either deletion of the last five residues (“del5”) or addition of an extra five C-terminal residues (-KASTN*; “add5”) resulted in dramatic stabilization in the context of the GPS system **(Fig. 2B)**. Next, to define the putative C-degron motif at amino acid resolution, we performed site-saturation mutagenesis of the ZMYND19 C-terminus to identify mutations that prevent degradation. Leveraging microarray oligonucleotide synthesis, we generated a ZMYND19 library in which each of the last 36 residues were mutated to all other possible amino acids and then measured the stability of the mutant pool by FACS followed by Illumina sequencing **(Fig. 2C)**. Stability profiles were highly concordant across two replicate experiments **(Fig. S3A and Table S2)**. These data highlighted the importance of the three C-terminal residues, -PER*, with the residues at the −2 and −3 positions appearing particularly critical for degron activity **(Fig. 2D)**. We validated this by assessing the stability of a panel of individual mutants, where addition of a single glutamic acid residue (“addD”), deletion of the last three residues (“del3”) or substitutions targeting any of the three terminal residues resulted in ZMYND19 stabilization **(Fig. 2E)**. The lack of effective commercial antibodies prevented us from directly monitoring the endogenous ZMYND19 protein. However, this effect was not limited to the GFP-fusion protein, as an HA-epitope tagged version of ZMYND19 was far more abundant upon deletion of the C-terminal residues as compared to the wild-type protein **(Fig. 2F)**. Furthermore, cycloheximide chase proteomics^32^ identifies ZMYND19 as one of the most unstable proteins in the human proteome across multiple cell lines, supporting the idea that the endogenous protein is regulated similarly. Altogether, these data indicate that ZMYND19 is an unstable protein due to a C-terminal degron.

**Figure 2.**
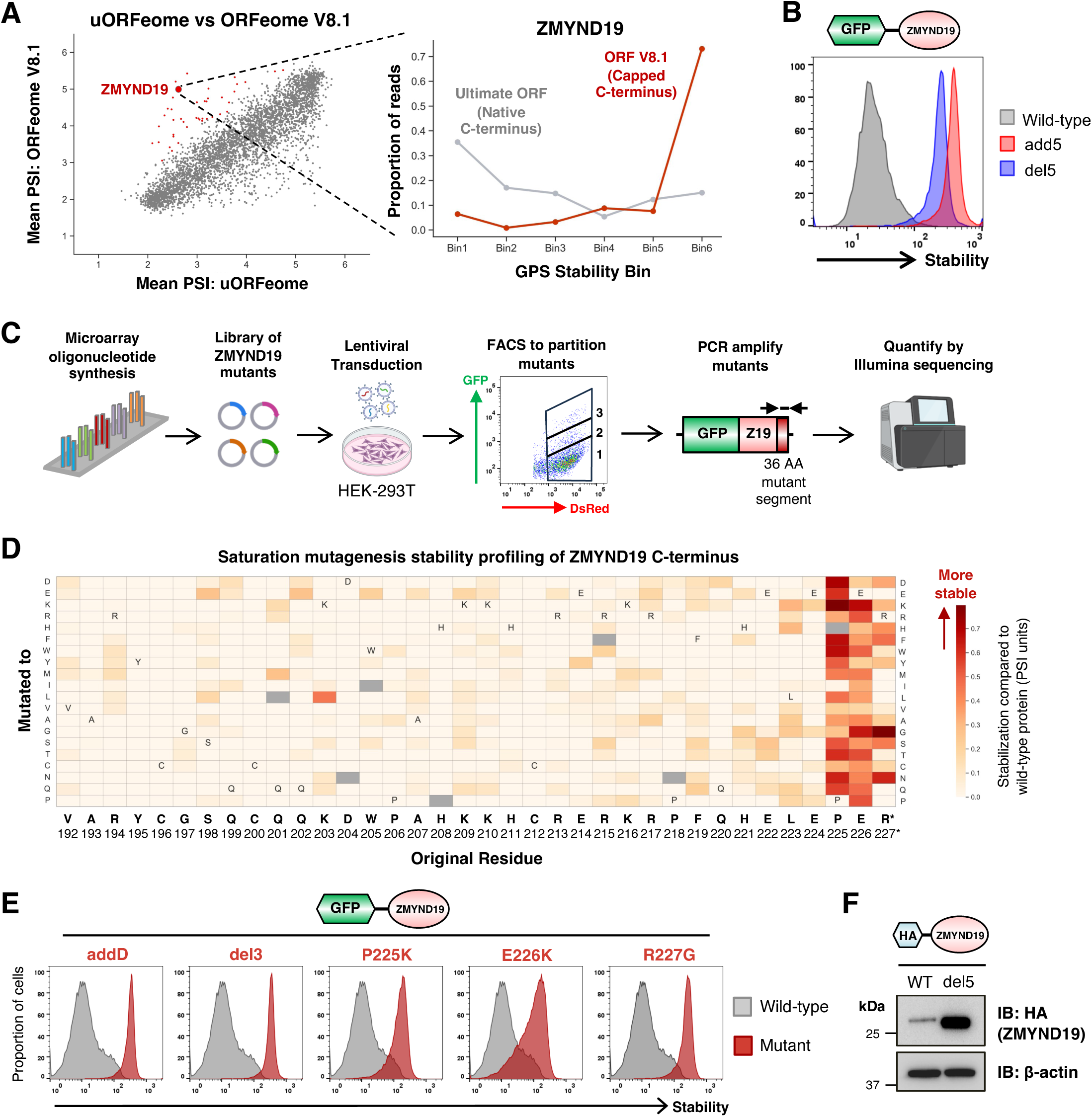
ZMYND19 harbors a C-terminal degron. **(A)** Comparative stability profiling identifies ZMYND19 as a protein harboring a candidate C-terminal degron, as the protein is much more stable when assayed in the context of the ORFeome V8.1 library with a capped C-terminus. **(B)** The C-terminus of ZMYND19 is responsibility for its instability, as both appending five additional C-terminal residues (-KASTN*; “add5”) or deleting five C-terminal residues (“del5”) resulted in stabilization. **(C-F)** Saturation mutagenesis of the ZMYND19 C-terminus defines a C-degron at amino acid resolution. **(C)** Schematic representation of the stability profiling experiment using a site-saturation mutagenesis library targeting the C-terminal 36 residues of ZMYND19. **(D)** Defining the ZMYND19 C-terminal degron. Each cell of the heatmap represents the stability of an individual substitution mutant; the darker the red color, the greater the degree of stabilization compared to the wild-type protein. **(E)** Individual validation experiments. Addition of a single aspartic acid residue (“addD”), deletion of three C-terminal residues (“del3”) or the indicated substitution mutations all resulted in ZMYND19 stabilization when assayed in the context of the GPS expression vector. **(F)** These effects were not limited to the GFP-fusion protein, as HA-tagged ZMYND19 was also more abundant upon deletion of five C-terminal residues (“del5”) as assessed by immunoblot.

### The CTLH E3 ligase complex targets ZMYND19 for proteasomal degradation

We next sought to identify the degradative machinery responsible for recognition of the C-terminal degron exposed by ZMYND19. Degradation of GPS-ZMYND19 was blocked by the proteasome inhibitor Bortezomib and the E1 inhibitor TAK-243, indicating ubiquitin-dependent proteasomal degradation **(Fig. S3B)**. To identify the E3 ligase responsible, we performed a genome-wide CRISPR/Cas9 genetic screen **(Fig. S3C)** which identified *MKLN1* (encoding Muskelin), *MAEA* and *TWA1* as significant hits **(Fig. 3A and Table S3)**. All three genes encode subunits of the CTLH (C-terminal to LisH) E3 ligase complex **(Fig. 3B)**. CTLH is a multi-subunit E3 ligase complex which is conserved among eukaryotes^33,34^. In yeast, the orthologous GID (glucose-induced degradation) complex plays an important role in gluconeogenesis by degrading several critical gluconeogenic enzymes in glucose-replete conditions. Intriguingly, substrate specificity is conferred via the Gid4 subunit, which targets N-terminal proline degrons^35–38^. The critical substrates of the CTLH complex in human cells remain incompletely defined^39^, but human GID4 also binds N-terminal proline^40,41^. Taken together with our findings, these data suggest that recognition of degrons located at protein termini may be central to substrate selection by CTLH.

**Figure 3.**
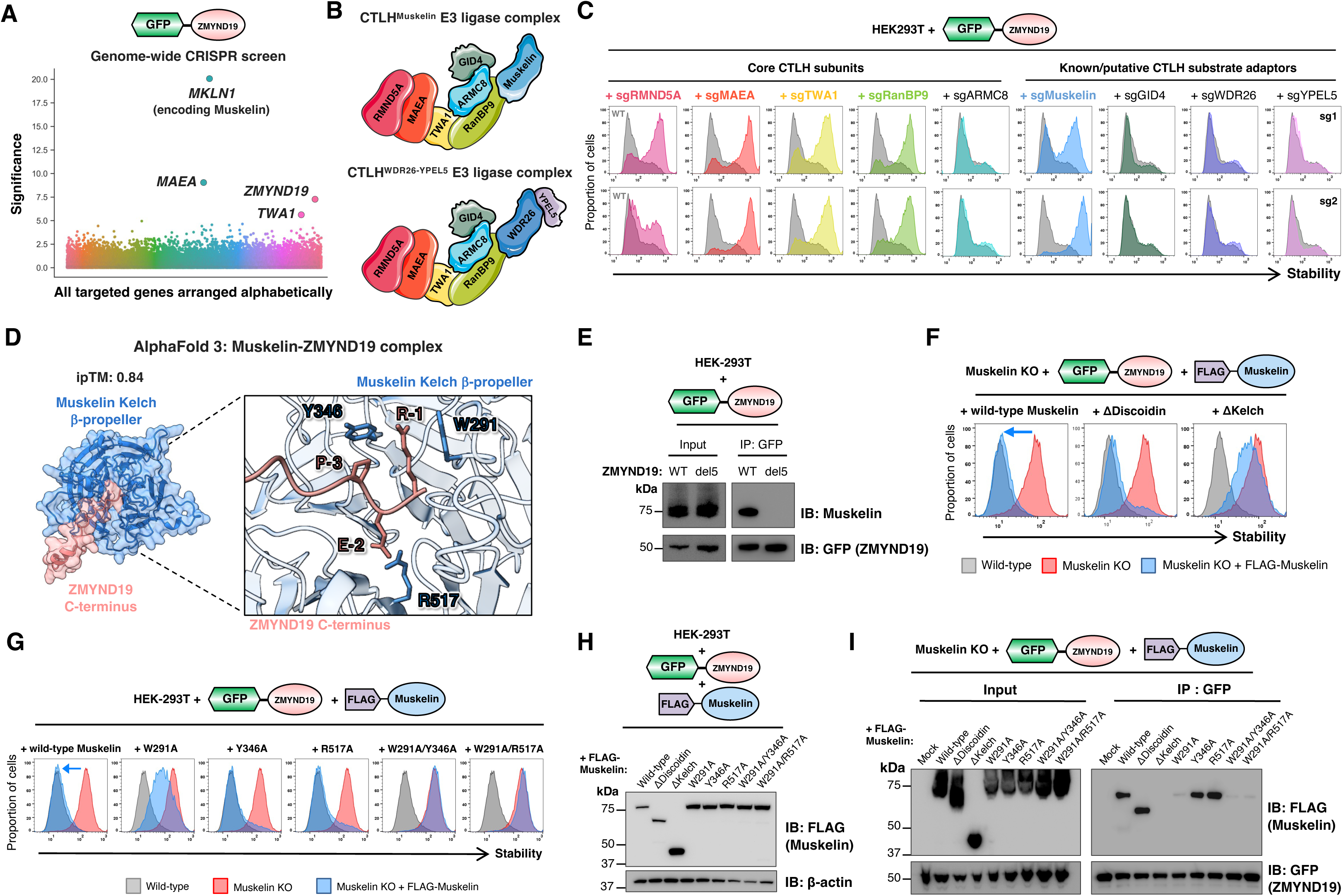
The Muskelin substrate adaptor of the CTLH E3 ligase complex targets the ZMYND19 C-terminal degron. **(A)** A genome-wide CRISPR screen identifies genes required for ZMYND19 degradation. All targeted genes are arranged alphabetically on the x-axis, with the −log10 of the MAGeCK “pos|score” metric reported on the y-axis. ZMYND19 itself also emerged as a significant hit alongside three CTLH components, presumably because CRISPR-mediated disruption of the exogenous GPS-ZMYND19 construct generates truncated GFP-fusion proteins lacking the C-terminal degron. **(B)** Schematic representation of the subunits of the CTLH E3 ligase complex. Muskelin and WDR26 are both orthologues of the single yeast protein Gid7 and are thought to exist in mutually exclusive CTLH complexes^77^. **(C)** Defining the CTLH components required for ZMYND19 degradation. Cells expressing GPS-ZMYND19 were transduced with Cas9 and sgRNAs targeting the indicated genes, and ZMYND19 stability assessed by flow cytometry seven days later. **(D)** AlphaFold 3 predicts an interaction between the β-propellor formed by the kelch repeats of Muskelin and the unstructured C-terminus of ZMYND19. **(E)** ZMYND19 binds Muskelin via its C-terminus. Muskelin co-immunoprecipitates with wild-type ZMYND19, but not with a ZMYND19 mutant lacking its last five residues (“del5”). **(F-I)** Defining the Muskelin residues required for recognition of the ZMYND19 C-terminal degron through genetic complementation. **(F)** ZMYND19 degradation in Muskelin KO cells can be restored by a Muskelin mutant lacking the discoidin domain, but not by a mutant lacking the kelch repeats. **(G)** Mutation of residues predicted to contact the ZMYND19 C-degron abolishes degradative activity. Individual mutation of W291 results in a Muskelin mutant that is only partially able to restore ZMYND19 degradation in Muskelin KO cells, whereas double mutants involving W291 and either Y346 or R517 abolish Muskelin activity. **(H)** Confirmation of expression of the mutant proteins by immunoblot. **(I)** Muskelin loss-of-function mutants fail to interact with ZMYND19. The interaction between the indicated FLAG-tagged Muskelin mutants and GFP-tagged ZMYND19 was assessed by co-immunoprecipitation. (In contrast to lysis in 1% SDS (H), Muskelin appears as a smear following lysis in 1% IGEPAL.)

We validated a role for the CTLH complex in ZMYND19 degradation through individual CRISPR/Cas9-mediated gene disruption experiments. With the exception of *ARMC8*, targeting any of the subunits considered part of the CTLH “core”– *RMND5A*, *MAEA*, *TWA1* or *RANBP9* – abolished ZMYND19 degradation **(Fig. 3C)**. Conversely, amongst the putative CTLH substrate adaptors, only ablation of *MKLN1* (encoding Muskelin) – but not *GID4*, *WDR26* or *YPEL5* – stabilized GPS-ZMYND19 **(Fig. 3C)**. To confirm that the lack of effect with these perturbations could not be explained by inefficient CRISPR-mediated knockout, we generated single cell knockout (KO) clones, verified successful gene disruption **(Fig. S3D-E)**, and then assessed the stability of GPS-ZMYND19. Unlike in MAEA, TWA1 and Muskelin KO cells, ZMYND19 degradation was unaffected in ARMC8, GID4, WDR26 and YPEL5 KO clones **(Fig. S3F)**. HA-tagged ZMYND19 was also far more abundant in MAEA KO and Muskelin KO cells than in wild-type cells **(Fig. S3G)**, and, further supporting the notion that the endogenous protein is regulated in the same way, proteomic datasets have identified ZMYND19 as a CTLH interacting partner^42^ and suggest increased ZMYND19 abundance upon loss of CTLH subunits^43,44^.

### The CTLH substrate adaptor Muskelin targets the ZMYND19 C-terminal degron

Our data thus far supported the hypothesis that the CTLH E3 ligase uses its substrate adaptor Muskelin to recruit ZMYND19 to the complex for ubiquitination and proteasomal degradation. This conclusion was supported by structural modelling, with AlphaFold 3 ^45^ predicting with high confidence (ipTM = 0.84) an interaction between the disordered C-terminus of ZMYND19 and the β-propellor formed by the kelch repeats of Muskelin **(Fig. 3D)**. We validated this model in several ways. First, we demonstrated that the interaction between ZMYND19 and Muskelin was dependent on the C-terminus of ZMYND19: immunoprecipitation of GFP-tagged ZMYND19 pulled down Muskelin, but a ZMYND19 mutant lacking the terminal five residues did not **(Fig. 3E)**. Second, we used genetic complementation experiments to identify the critical domain(s) of Muskelin required for ZMYND19 degradation. Exogenous expression of either wild-type Muskelin or a Muskelin mutant lacking the N-terminal discoidin domain fully restored GPS-ZMYND19 degradation in Muskelin knockout cells, whereas a Muskelin mutant lacking the kelch repeats did not **(Fig. 3F)**. Third, we further exploited this assay to examine the effect of point mutations that, based on the AlphaFold 3 structural model, would be predicted to impair ZMYND19 C-degron recognition. While individual substitutions had limited impact on Muskelin function, with mutation of W291 (predicted to contact the R-1 residue of ZMYND19) partially abrogating ZMYND19 degradation and mutation of Y346 (predicted to contact the P-3 residue) and R517 (predicted to contact the E-2 residue) having no detrimental effect, we found that double mutants were completely non-functional **(Fig. 3G-H)**. These effects were indeed due to an inability of the Muskelin mutants to interact with ZMYND19, as assessed by co-immunoprecipitation experiments **(Fig. 3I)**. Thus, genetic interrogation of the AlphaFold 3 model supported the notion that the β-propellor formed by the kelch repeats of Muskelin mediate selective recognition of the ZMYND19 C-terminal degron.

### Cell-to-cell variability in ZMYND19 stability reveals conditional regulation of CTLH activity

We observed cell-to-cell variability in the stability of the GFP-ZMYND19 fusion construct **(Fig. 4A)**. GFP fluorescence was typically undetectable in >90% of the cells, indicative of very efficient ZMYND19 degradation, but remained bright in ∼5-10% of the cells, suggesting conditional inactivation of CTLH-mediated ZMYND19 degradation **(Fig. 4A)**. The proportion of the cells in which ZMYND19 was stabilized was variable, however, and, on occasion, rose towards 50% without obvious changes in the cell culture conditions **(Fig. 4A)**. Moreover, following isolation of the GFP^bright^ population by FACS, the population quickly reverted towards a GFP^dim^ state **(Fig. 4B)**.

**Figure 4.**
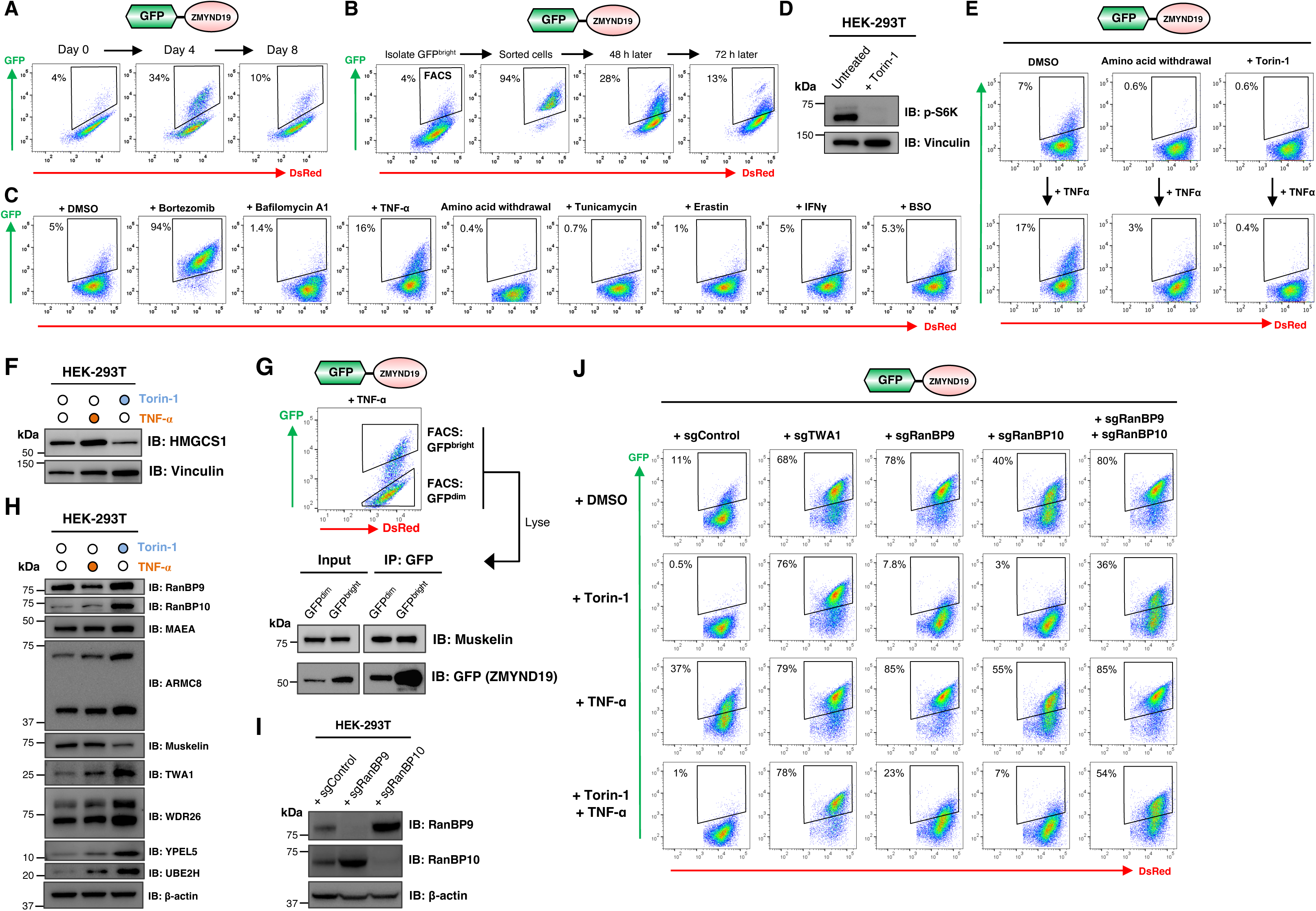
Cell-to-cell variability in ZMYND19 stability reveals conditional regulation of CTLH activity. **(A-B)** Cell-to-cell variability in ZMYND19 stability. **(A)** ZMYND19 expressed in the context of the GPS system escapes degradation in a variable proportion of the cells. **(B)** Following isolation of the GFP^bright^ cells by FACS, the population quickly reverted towards the GFP^dim^ state. **(C)** A chemical screen to identify conditions which impact ZMYND19 degradation. HEK-293T cells expressing ZMYND19 in the context of the GPS vector were subjected to the indicated treatments and the effect on ZMYND19 degradation assessed by flow cytometry. **(D-E)** Inhibition of mTOR signaling potentiates ZMYND19 degradation. **(D)** Overnight treatment with the mTOR inhibitor Torin-1 abolishes phosphorylation of p70 S6 kinase. **(E)** Amino acid withdrawal or Torin-1 treatment enhances ZMYND19 degradation, overriding the effects of TNF-α stimulation. See also Fig. S4C. **(F)** Conditional regulation of the CTLH^GID4^ substrate HMGCS1. HMGCS1 is more abundant following TNF-α stimulation and less abundant following Torin-1 treatment, as assessed by immunoblot. **(G)** The association between ZMYND19 and Muskelin is maintained in GFP^bright^ cells where GPS-ZMYND19 escapes degradation. Following overnight TNF-α stimulation, cells expressing GPS-ZMYND19 were partitioned into GFP^dim^ and GFP^bright^ populations by FACS and the interaction between ZMYND19 and Muskelin assessed via co-immunoprecipitation. **(H)** Effect of TNF-α stimulation and mTOR inhibition on the abundance of CTLH subunits, as assessed by immunoblot. **(I-J)** Differential requirements for RANBP9 and RANBP10 for CTLH-mediated degradation of ZMYND19 upon mTOR inhibition. Using sgRNAs that result in efficient gene disruption as assessed by immunoblot **(I)**, the effects of targeting RANBP9 and RANBP10 either individually or simultaneously on ZMYND19 degradation were assessed by flow cytometry **(J)**. Ablation of RANBP9 prevents ZMYND19 degradation at steady state; upon mTOR inhibition, however, targeting of RANBP9 has little impact on ZMYND19 stability, and dual ablation of RANBP9 and RANBP10 is necessary before abrogated degradation is observed.

To examine potential stimuli which might result in conditional regulation of the CTLH-mediated degradation of ZMYND19, we subjected cells expressing GPS-ZMYND19 to a broad range of insults and measured the proportion of GFP^+^ cells by flow cytometry. These experiments revealed that stimulation with the inflammatory cytokine tumor necrosis factor alpha (TNF-α) reliably increased the proportion of cells in which GPS-ZMYND19 was stable **(Fig. 4C and Fig. S4A)**. TNF-α treatment did not seem to generally inhibit proteasomal degradation, as the stability of GPS-CHGA, for example, a substrate of Cul2^KLHDC3^ **(Fig. 1G)**, was unaffected **(Fig. S4B)**. Unexpectedly, we also noted that amino acid starvation decreased the proportion of cells in which GPS-ZMYND19 was stable **(Fig. 4C and Fig. S4A)**. This effect was particularly striking when combined with TNF-α treatment, where amino acid starvation exerted a dominant effect and kept GPS-ZMYND19 unstable in almost 100% of the cells **(Fig. 4E and Fig. S4C)**. Further probing the mechanism through which amino acid deprivation might stimulate the degradative activity of CTLH, we considered a possible role for mTOR as a key sensor of amino acid sufficiency^46^. Indeed, inhibiting mTOR signaling with the small molecule inhibitor Torin-1^47^ phenocopied these effects **(Fig. 4D-E and Fig. S4C)**. Thus, we concluded that ZMYND19 degradation is conditionally regulated: TNF-α stimulation favors CTLH inactivation resulting in ZMYND19 stabilization, while amino acid deprivation upstream of loss of mTOR activity potentiates CTLH-mediated ZMYND19 degradation.

During the preparation of this manuscript, mTOR inhibition was reported to stimulate the degradation of HMGCS1 by CTLH^GID4^ and UCK2 by CTLH^WDR26^ ^44,48^. Together with the knowledge that mTOR inhibition stimulates the degradation of ZMYND19 by CTLH^Muskelin^, these data suggest that loss of mTOR signaling acts as a general mechanism to potentiate CTLH activity irrespective of the mode of substrate recruitment. Similarly, the inhibitory effects of TNF-α stimulation on CTLH also appeared to function independently of the mechanism of substrate recruitment, as TNF-α also increased the abundance of HMGCS1 as measured by immunoblot **(Fig. 4F)**, and, after isolating the GFP^bright^ population in which ZMYND19 degradation was abrogated by FACS, we still observed Muskelin efficiently co-immunoprecipitating with its substrate **(Fig. 4G)**.

Determining the mechanistic basis underpinning the inhibitory effects of TNF-α signaling and the stimulatory effects of mTOR inhibition on CTLH function remains a key goal for future study. We found that neither mTOR inhibition nor TNF-α stimulation resulted in dramatic changes in CTLH complex subunit composition **(Fig. S4D)**, although we did note some changes in the abundance of CTLH subunits **(Fig. 4H)**. However, we did find some support for the notion that alternative paralog usage might play a role^49^. Although RANBP9 is required for the degradation of GPS-ZMYND19 at steady-state, surprisingly we found that it was dispensable for the CTLH-mediated degradation of GPS-ZYMND19 following mTOR inhibition **(Fig. 4I-J)**. CRISPR/Cas9-mediated disruption of the RANBP9 paralog RANBP10 also did not prevent ZMYND19 degradation upon mTOR inhibition, with simultaneous targeting of both RANBP9 and RANBP10 required to abrogate this effect **(Fig. 4I-J)**. Thus, mTOR inhibition may alter the properties of both RANBP9 and RANBP10 which leads to potentiation of CTLH activity towards its substrates. Overall, these data reveal that the CTLH E3 ligase complex is subject to conditional regulation, with TNF-α stimulation impairing CTLH activity and mTOR inhibition enhancing it, and, while these effects may involve alterations to the behavior of the paralogous subunits RANBP9 and RANBP10, the molecular mechanism(s) remains to be defined.

### Parallel genetic and proteomic approaches identify additional substrates of the CTLH^Muskelin^ C-degron pathway

Next we sought to leverage the insight that mTOR inhibition potentiates CTLH activity to identify additional substrates targeted via the CTLH^Muskelin^ C-degron pathway. We began by performing further stability profiling screens using the GPS-uORFeome library, comparing wild-type cells versus MAEA KO and TWA1 KO cells following treatment with Torin-1 **(Fig. 5A)**. This experiment identified four high-confidence CTLH substrates: ZMYND19, which served as a positive control, plus three poorly characterized proteins: AAMP (angio-associated migratory protein), AEN (apoptosis-enhancing nuclease) and NATD1 (N-acetyltransferase domain containing 1) **(Fig. 5B, Fig. S5A and Table S4)**. Individual validation experiments revealed that, like ZMYND19, AAMP and AEN were both substrates of CTLH^Muskelin^, whereas NATD1 degradation was independent of Muskelin but required both the WDR26 and YPEL5 substrate adaptors **(Fig. 5C and Fig. S5B)**. In parallel, we also employed a proteomic approach. Following CRISPR/Cas9-mediated gene disruption in cells expressing GPS-ZMYND19, we purified populations of knockout cells lacking either the core CTLH subunit TWA1 or the substrate adaptor Muskelin by FACS **(Fig. S5C)** and performed tandem mass tag (TMT) proteomics **(Fig. 5D)**. This revealed a handful of proteins exhibiting concordantly increased abundance to a greater extent than the positive control substrate HMGCS1, including AAMP **(Fig. 5E-F and Table S5)**.

**Figure 5.**
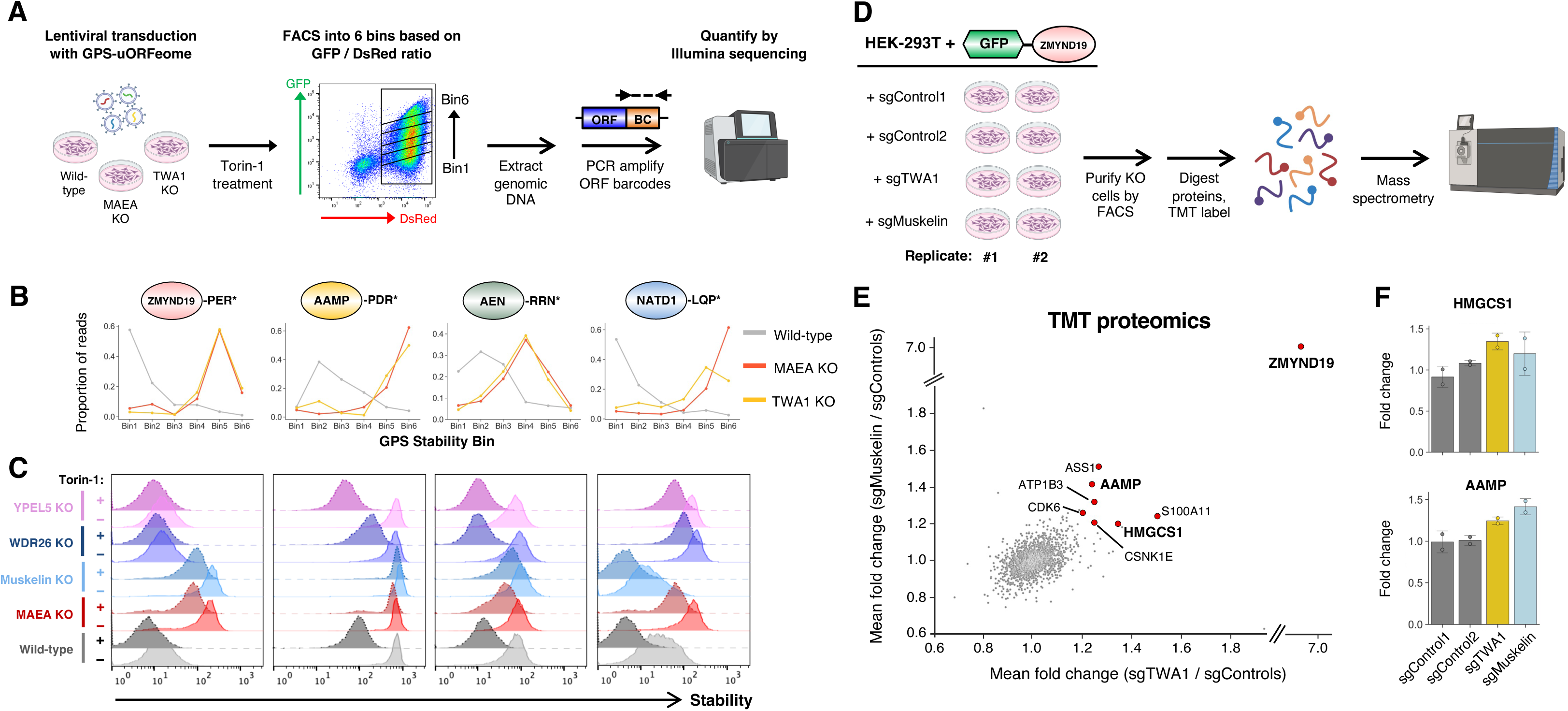
Parallel genetic and proteomic screens identify additional substrates of the CTLH^Muskelin^ C-degron pathway. **(A)** Schematic representation of the stability profiling experiment, in which the Ultimate ORFeome GPS expression library was used to identify proteins exhibiting increased stability in MAEA KO and TWA1 KO cells. **(B-C)** Identification of AAMP, AEN and NATD1 as CTLH substrates. Screen profiles are depicted in **(B)**, while individual validation experiments by flow cytometry in the presence and absence of Torin-1 (dotted lines) are shown in **(C)**. See also Fig. S5B. **(D)** Schematic representation of the proteomic experiment, in which TMT mass spectrometry was used to identify proteins more abundant in cells lacking the CTLH subunits TWA1 and Muskelin. Cells expressing GPS-ZMYND19 were used to allow purification of populations of cells in which TWA1 and Muskelin had been successfully ablated by FACS. See also Fig. S5C. **(E-F)** AAMP is more abundant in cells lacking CTLH subunits. The scatterplot in **(E)** represents the mean fold change of each protein quantified in both TWA1 KO and Muskelin KO cells, with proteins showing a >1.2-fold increase in abundance in both conditions highlighted in red; the performance of AAMP compared to the known substrate HMGCS1 is depicted in **(F)**. Exogenous expression of ZMYND19 precludes assessment of the true magnitude of the stabilization of the endogenous ZMYND19 protein in this experiment.

AAMP stood out as an intriguing hit: the protein was identified by both genetic and proteomic screens, and, strikingly, terminates with the amino acid sequence -PDR*, a very similar motif to the ZMYND19 -PER* C-terminal degron targeted by Muskelin. Furthermore, although its function is uncharacterized, sequence analysis using Foldseek^50^ revealed extensive structural homology with yeast Sqt1, a ribosome biogenesis factor **(Fig. S5D)**. AEN, on the other hand, does not terminate with a candidate Muskelin C-degron motif, but multiple large-scale interactome studies^51^ suggest that it interacts with AAMP, raising the possibility that AEN may be recruited to CTLH via AAMP. The function of AEN is also unknown, but, alongside ISG20 and ISG20L2, it constitutes a member of the ISG20 family of exonucleases **(Fig. S5E)**. ISG20 expression is induced by interferon signaling and is thought to function in antiviral defense^52^, while ISG20L2 is an essential protein whose exoribonuclease activity is involved in the maturation of both 5.8S and 18S pre-ribosomal RNA^53,54^.

### The interacting partners AAMP and AEN are both substrates of the CTLH^Muskelin^ C-degron pathway

We began by investigating the effect of manipulating the C-terminus of AAMP. Unlike the wild-type protein, AAMP lacking its terminal five residues (“del5”) was not destabilized upon mTOR inhibition **(Fig. 6A)** and did not associate with Muskelin **(Fig. 6B)**. Endogenous AAMP also interacted with Muskelin **(Fig. S5F)**, and its abundance was increased in Muskelin KO cells **(Fig. S5G)**. AlphaFold 3 predicts an interaction between the unstructured C-terminus of AAMP and the β-propellor formed by the kelch repeats of Muskelin **(Fig. 6C)** which mirrors that of the predicted ZMYND19-Muskelin interaction **(Fig. S5H)**. Furthermore, Muskelin mutants affecting the predicted binding interface were unable to bind AAMP **(Fig. S5I-J)** and did not support AAMP degradation upon mTOR inhibition **(Fig. 6D)**. Thus, AAMP is a substrate of the C-degron pathway regulated by CTLH^Muskelin^.

**Figure 6.**
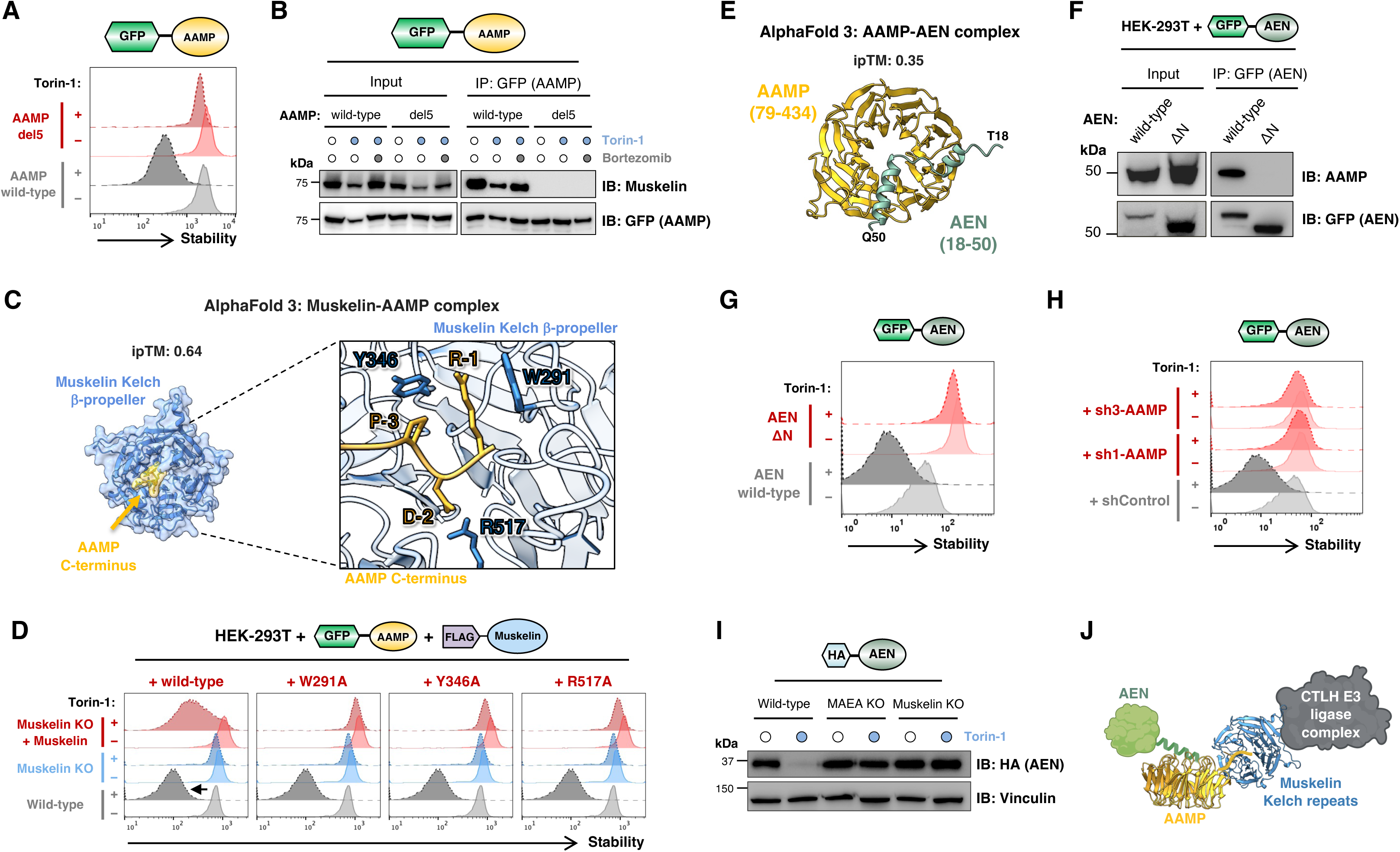
The CTLH^Muskelin^ C-degron pathway regulates the stability of AAMP and its binding partner AEN. **(A-D)** AAMP exposes a C-degron targeted by Muskelin. **(A)** Wild-type AAMP expressed in the context of the GPS system is degraded upon treatment with Torin-1, but not upon removal of five C-terminal residues (“del5”). **(B)** Wild-type AAMP interacts with Muskelin as assessed through co-immunoprecipitation experiments, but not following deletion of its C-terminus. **(C)** AlphaFold 3 model of the AAMP-Muskelin interaction. **(D)** Muskelin mutants targeting the predicted AAMP binding interface are non-functional. Unlike wild-type Muskelin, exogenous expression of Muskelin variants harboring mutations in residues predicted to contact the AAMP C-terminus are incapable of restoring AAMP degradation following mTOR inhibition. **(E-G)** AEN interacts with AAMP via its N-terminus. **(E)** AlphaFold 3 predicts an interaction between an N-terminal a-helix of AEN (green) and the β-propellor formed by the WD40 repeats of AAMP (gold). **(F)** Wild-type AEN associates with AAMP as assessed by co-immunoprecipitation experiments, but not an AEN mutant lacking its N-terminus (“ΔN”) does not; accordingly, wild-type AEN but not the ΔN mutant is destabilized upon Torin-1 treatment **(G)**. **(H)** AEN is subject to collateral degradation through the CTLH^Muskelin^ C-degron pathway via AAMP. Cells expressing AEN in the context of the GPS vector were transduced with shRNAs targeting AAMP and AEN stability was monitored by flow cytometry following overnight treatment with Torin-1. The destabilization of AEN induced by mTOR inhibition is abolished in cells depleted of AAMP. **(I)** The abundance of HA-tagged AEN is decreased following treatment with Torin-1 in wild-type cells, but not in cells lacking the CTLH subunits MAEA or Muskelin. **(J)** Model summarizing the collateral degradation of the AAMP binding partner AEN via the CTLH^Muskelin^ C-degron pathway.

We next considered the fate of AEN, which was also robustly degraded by CTLH^Muskelin^ upon mTOR inhibition **(Fig. 5C)**. AlphaFold 3 proposes that the N-terminal α-helix of AEN may be accommodated by the β-propellor formed by the WD40 repeats of AAMP, albeit at modest confidence (ipTM = 0.35) **(Fig. 6E)**. Supporting this model, immunoprecipitation of wild-type AEN pulled-down AAMP, whereas an AEN mutant lacking the N-terminal helix (“ΔN”) did not **(Fig. 6F)**. Furthermore, the AEN ΔN mutant was refractory to Torin-1-induced degradation **(Fig. 6G)**. To directly test if AAMP is required to bridge the interaction between AEN and CTLH, we depleted cells of AAMP **(Fig. S6A)** and assessed the effect on AEN stability. We found that AAMP knockdown abolished the degradation of GPS-AEN induced upon Torin-1 treatment **(Fig. 6H)**. Moreover, the interaction between AEN and Muskelin was preserved upon reconstitution of AAMP knockdown cells with shRNA-resistant full-length AAMP but not following reconstitution with AAMP mutants bearing altered C-termini **(Fig. S6B-C)**, and AEN was efficiently degraded upon mTOR inhibition only when AAMP with an intact C-terminus was present **(Fig. S6D)**. The lack of effective commercial antibodies precluded us from assessing whether endogenous AEN behaved similarly. However, AEN fused to a small, lysine-less HA epitope tag exhibited decreased abundance upon mTOR inhibition in a manner that depended on CTLH **(Fig. 6I)**, and we also note that AEN emerged as a leading candidate CTLH substrate in a recent proteomic study^44^. Altogether, these data demonstrate that AEN is subject to collateral degradation through the CTLH^Muskelin^ C-degron pathway via AAMP **(Fig. 6J)**.

We also considered the fate of ISG20L2, which shares 49% identity with AEN and is predicted to possess an N-terminal α-helix with similar characteristics. We confirmed that ISG20L2 also interacts with AAMP via a co-immunoprecipitation assay **(Fig. S6E)**, and AlphaFold 3 suggests that the N-terminal α-helix of ISG20L2 could interact with AAMP in an analogous manner to that of AEN **(Fig. S6F)**. Unlike AEN, however, ISG20L2 did not appear to be a substrate of collateral degradation involving CTLH^Muskelin^, as we observed no difference in the abundance of ISG20L2 following mTOR inhibition or ablation of Muskelin **(Fig. S6G-H)**. We conclude that although both AEN and ISG20L2 associate with AAMP, only AEN is targeted for proteasomal degradation by CTLH^Muskelin^.

### AAMP is required for ribosome biogenesis in human cells

Finally, we sought to explore the physiological role of AAMP in ribosome biogenesis. In *Saccharomyces cerevisiae*, the AAMP ortholog Sqt1 is well-characterized as a chaperone for the large ribosomal subunit uL16 (formerly designated as Rpl10)^55–57^. uL16 is one of the last subunits to be incorporated into pre-60S particles in the cytosol^56^, triggering the release of the final biogenesis factors to complete the catalytic center of the ribosome and license subunit joining to form translationally competent 80S ribosomes^58^. Sqt1 binds uL16 co-translationally, shielding a positively charged α-helix that will later associate with helix H89 of the 25S rRNA^57^. Complemented by data from large-scale interactome studies^59^ which suggest an association between AAMP and uL16 in human cells, we therefore tested the hypothesis that AAMP is required for mammalian ribosome biogenesis.

We first confirmed that AAMP binds uL16 through co-immunoprecipitation experiments **(Fig. 7A)**. The crystal structure^60^ of Sqt1 in complex with yeast uL16 demonstrates that the β-propellor formed by the WD40 repeats of Sqt1 recognizes an α-helix located at the N-terminus of uL16 **(Fig. 7B)**, and AlphaFold 3 predicts with high confidence (ipTM = 0.87) that AAMP engages human uL16 in an analogous manner **(Fig. 7C)**. In support of this model, a uL16 mutant lacking its N-terminal helix failed to associate with AAMP **(Fig. 7D)**; furthermore, fusing just the N-terminus of uL16 (residues 1-28) to GFP was sufficient to mediate destabilization upon mTOR inhibition **(Fig. 7E, right panel)**. Full-length uL16 did not appear to be a substrate of CTLH-mediated degradation via AAMP, however **(Fig. 7E, left panel)**, and endogenous uL16 levels were unaffected in the absence of CTLH subunits **(Fig. 7F)**.

**Figure 7.**
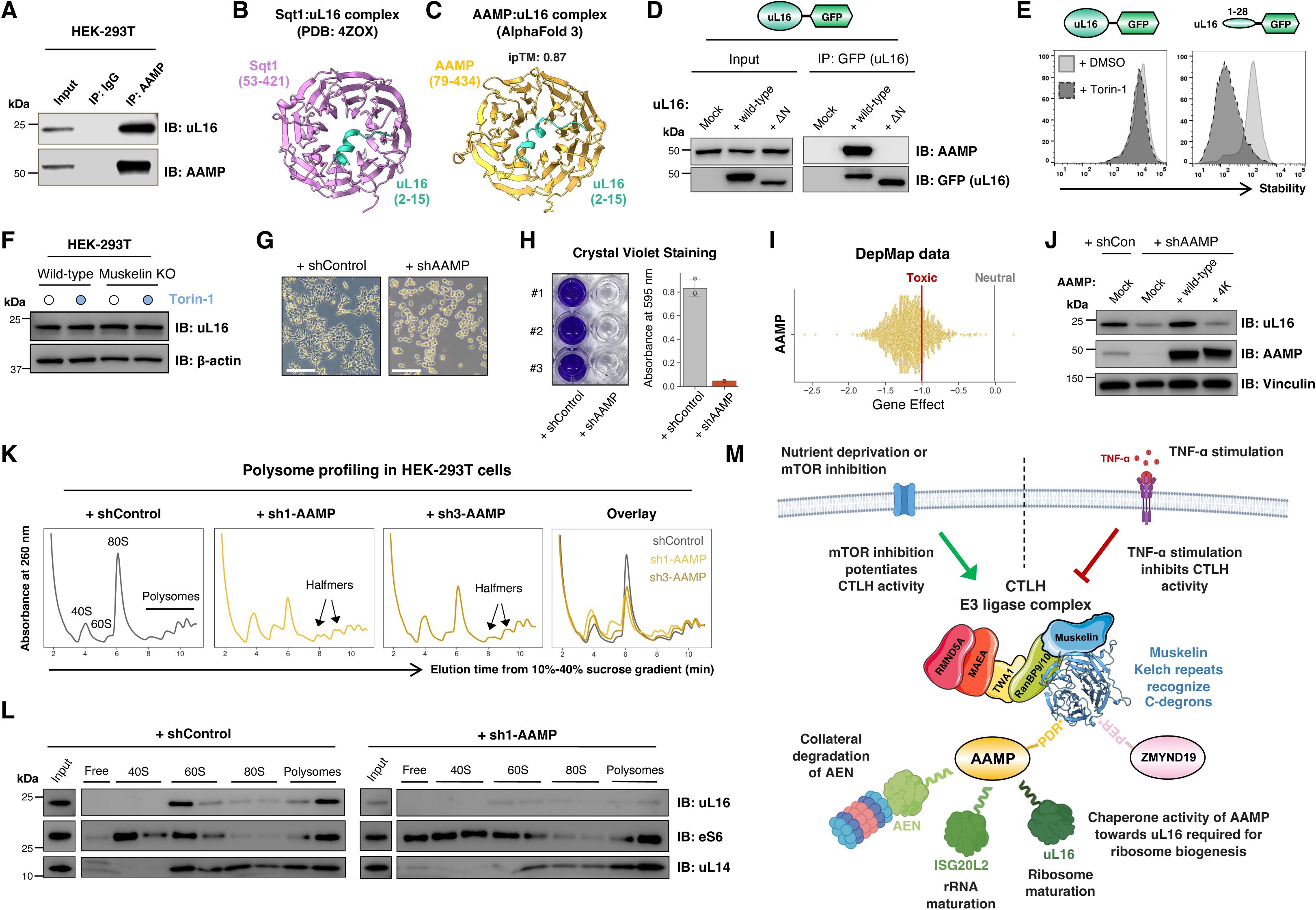
AAMP is the ortholog of yeast Sqt1 and is essential for ribosome biogenesis in human cells. **(A-D)** AAMP associates with the N-terminus of uL16. **(A)** uL16 co-immunoprecipitates with AAMP. **(B)** The crystal structure of Sqt1 in complex with yeast uL16 reveals a basic alpha-helix at the N-terminus of uL16 accommodated in an acidic pocket formed by the WD40 repeats of Sqt1; AlphaFold 3 predicts that human uL16 engages AAMP in an analogous manner **(C)**, and deletion of the N-terminus of uL16 abolishes its association with AAMP **(D)**. **(E-F)** uL16 is not a substrate of collateral degradation by CTLH^Muskelin^ via AAMP. Although a construct comprising the first 28 residues of uL16 fused to GFP was subject to degradation upon Torin-1 treatment, full-length uL16 fused to GFP was not **(E)**. Furthermore, the abundance of endogenous uL16 was not affected by Torin-1 treatment or loss of Muskelin **(F)**. **(G-J)** *AAMP* is an essential gene. Following transduction with an shRNA expression vector targeting AAMP, HEK-293T cells exhibited a rounded morphology at day 5 post-transduction (scale bar, 100 µm) **(G)** and had died by day 7 post-transduction as assessed by crystal violet staining **(H)**. Furthermore, CRISPR screen data from the DepMap project demonstrates that disruption of AAMP is highly detrimental to cell viability across >1000 cell lines **(I)**. The DepMap “Gene Effect” score^78^ is scaled such that the effect size for nonessential genes is centered at 0 (gray line), and the median effect of a group of known essential genes is centered at −1 (red line). **(J)** AAMP depletion decreases the protein abundance of uL16 as assessed by immunoblot; this effect could be rescued by expression of shRNA-resistant wild-type AAMP, but not by an AAMP charge-swap mutant (“4K”) unable to bind uL16. **(K-L)** AAMP depletion impairs ribosome maturation. Polysome profiling in AAMP knockdown cells **(K)** reveals an increase in the size of the 40S peak, a decrease in the size of the 60S and 80S peaks, and the appearance of “halfmers” in the polysome fraction. The distribution of uL16, eS6 and uL14 across fractions corresponding to the indicated peaks is shown in **(L)**. **(M)** Schematic depiction of the role of the CTLH^Muskelin^ C-degron pathway in regulating ribosome biogenesis via AAMP.

We went on to examine the functional consequences of AAMP ablation. In support of an indispensable role for AAMP in ribosome biogenesis, we found that loss of AAMP was extremely toxic **(Fig. 7G-H)**. Concordantly, data from the DepMap project reveals that CRISPR-mediated disruption of *AAMP* is uniformly deleterious across hundreds of human cell lines **(Fig. 7I)**. Consistent with a requirement for AAMP for uL16 function, we observed a substantial reduction in uL16 protein levels upon AAMP knockdown by immunoblot **(Fig. 7J)**. This effect could be rescued upon expression of shRNA-resistant wild-type AAMP, but not by a charge-swap (“4K”) mutant in which four acidic residues predicted to contact uL16 were mutated to lysine **(Fig. S7A-D)**. Intriguingly we also noted a similar effect on ISG20L2 abundance **(Fig. S7E)**, suggesting that AAMP may be required for the normal function of both uL16 and ISG20L2. Lastly, we observed significant defects in ribosome maturation upon AAMP depletion as assessed by polysome profiling **(Fig. 7K-L)**. Depletion of AAMP resulted in an increase in the size of the 40S peak, substantial reductions in both the 60S and 80S peaks and the appearance of “halfmers” in the polysome fractions, features consistent with a ribosomal subunit joining defect resulting from uL16 insufficiency **(Fig. 7K-L)**. Overall, these data suggest a model whereby the increased activity of CTLH following loss of mTOR activity downstream of nutrient deprivation could negatively regulate ribosome biogenesis through AAMP degradation via the CTLH^Muskelin^ C-degron pathway **(Fig. 7M)**.

## DISCUSSION

With the goal of delineating the physiological context within which C-degron pathways act, here we exploited two distinct human ORFeome libraries with differing C-terminal architectures to identify full-length proteins harboring C-degrons. Comparative stability profiling assessed the impact of manipulating the C-termini of thousands of human proteins, identifying ∼50 substrates bearing candidate C-degron motifs. Characterizing the mode of degradation of one of the leading candidates, ZMYND19, we uncovered a C-degron pathway regulated by the Muskelin substrate adaptor of CTLH E3 ligase complex which is under conditional control. We elucidated the function of one of the substrates of this pathway, AAMP, which serves an essential role in ribosome maturation via its chaperone activity towards uL16. Altogether, these data define a C-degron pathway which connects CTLH^Muskelin^ to the regulation of ribosome biogenesis.

Our comparative stability profiling screens concordantly identified ∼50 full-length proteins exhibiting substantially increased stability upon addition of extra residues at their C-terminus. Although this likely represents an underestimate of the total number of proteins regulated by C-degrons, particularly as the scale of our approach was limited to the ∼5000 proteins quantified across both libraries, further analysis of these substrates has the potential to identify many more C-degron pathways. Indeed, sequence analysis reveals that most of the candidate substrates do not bear known C-degron motifs, suggesting the existence of additional E3 ligases targeting protein C-termini. We also note that ∼100 proteins exhibited concordant destabilization upon C-terminal “capping” **(Fig. 1C)**, and thus we speculate that classes of protein other than E3 ubiquitin ligases may recognize “stabilon” motifs at protein termini to protect substrates from degradation. Although beyond the scope of this study, genetic interrogation of these proteins using the techniques employed here should reveal the molecular mechanisms involved.

Our follow-up work focused on ZMYND19 revealed that the CTLH E3 ligase complex regulates a C-degron pathway via its substrate adaptor Muskelin. An emerging theme amongst E3 ligases directed towards protein termini is the use of tandem repeat domains to achieve specificity, such as the ankyrin repeats of FEM1B^22^, the WD40 repeats of DCAF12^61^ and the tetratricopeptide repeats of APPBP2^62^. Muskelin is no exception, using the β-propellor architecture formed by its kelch repeats to engage the C-termini of its substrates. Our saturation mutagenesis of the ZMYND19 C-terminus highlighted a degron motif comprising either proline or isoleucine at the −3 position of the substrate, aspartic acid or glutamic acid at the −2 position, and a variety of candidate residues in addition to arginine at the terminal position **(Fig. 2D)**. One major advantage of high-resolution mapping of degron motifs is that it allows for bioinformatic prediction of additional substrates. Indeed, recent proteomic data^44^ identifies NAPL12 (terminating -IDR*) as one of the proteins whose abundance is greatly increased in both MAEA and Muskelin KO cells **(Fig. S8)**, suggesting that it is likely to represent an additional substrate of the Muskelin C-degron pathway identified here.

The substrate repertoire of the CTLH^Muskelin^ C-degron pathway could well be broader than our genetic data suggests, however. UNG2 was recently identified as a substrate of collateral degradation by CTLH^Muskelin^ via an intermediate protein FAM72A^63^; although the yippee-like domain of FAM72A also plays an important part in the interaction with Muskelin, the cryo-electron microscopy structure of the Muskelin-FAM72A-UNG2 complex shows the C-terminus of FAM72A accommodated by the kelch repeats of Muskelin^63^. However, the C-terminal sequence of FAM72A (-CIR*) does not closely resemble the -PER* or -PDR* motifs of ZMYND19 and AAMP respectively. Although the saturation mutagenesis technique is a powerful approach to define degron motifs at amino acid resolution, the results obtained may be specific to the context in which the screen was performed; thus, further genetic analysis of C-degrons targeted by Muskelin beyond ZMYND19 may be required to define the full repertoire of candidate degron motifs.

While the GID complex plays a well-understood metabolic role in the regulation of gluconeogenesis in yeast, the function of the orthologous CTLH complex in mammalian cells remains less clear. An important breakthrough is the discovery that CTLH activity is conditionally regulated. Our data supports the recent finding that loss of mTOR signaling potentiates CTLH activity^44^, and further extends this notion by demonstrating that TNF-α stimulation has the opposite effect and abrogates CTLH-mediated degradative activity. Accelerated degradation of HMGCS1 ^44^, the first enzyme in the mevalonate synthesis pathway, UCK2 ^48^, the rate-limiting enzyme in the pyrimidine salvage pathway, and AAMP, which we define here as an essential factor required for ribosome biogenesis, suggest that CTLH is likely to be an important effector downstream of mTOR, whereby enhanced degradation of a variety of substrates contributes towards the cell’s effort to mitigate the effects of nutrient deprivation. Equally, our observation that TNF-α stimulation impairs CTLH activity suggests that there must also be physiological circumstances where the opposite effect is required. Loss of CTLH subunits was recently reported to impair the growth of the intracellular bacteria in macrophages, suggesting that at steady-state CTLH activity restrains host anti-microbial responses^64^. Thus, it is tempting to speculate that one effect of pro-inflammatory TNF-α signaling might be to impair CTLH function and thus allow a rapid induction of innate immune pathways. TNF-α stimulation is unlikely to represent the optimal route to suppress CTLH activity, however, particularly as we only tested the effect of a handful of stressors and the inhibitory effects we observed were only partial **(Fig. 4 and Fig. S4)**. Moreover, TNF-α stimulation cannot account for the small proportion of cells in which ZMYND19 degradation is inhibited at steady-state **(Fig. 4A)**; thus, there must be additional pathways through which CTLH activity can be suppressed.

A priority for future work will be to define the molecular mechanisms underpinning the differential effects of TNF-α stimulation and mTOR inhibition on CTLH activity. One possibility is that these effects are direct, with for example the kinase activity of mTOR itself serving to phosphorylate a core CTLH subunit to tune the efficiency CTLH-mediated ubiquitination, but they could also occur via intermediate proteins and hence may not necessarily require post-translational modification of CTLH subunits. Given that these stimuli appear to affect all substrates examined, irrespective of their mode of recruitment to CTLH, our data supports the notion that these both stimuli are likely to impact the ability of the CTLH “core” to mediate the ubiquitination and degradation of these substrates. Interestingly, however, different substrates are affected to different extents: GPS-ZMYND19, for example, was degraded very efficiently at steady-state and hence exhibited little scope for further potentiation of degradation upon mTOR inhibition, whereas GPS-AAMP was essentially refractory to CTLH-mediated degradation until mTOR signaling was inhibited **(Fig. 5C)**. Determining the factors that underlie these contrasting effects may help illuminate the mechanistic basis for differential CTLH activity, and the genetic tools described here should serve to complement biochemical and structural approaches towards this goal.

Finally, our investigation into the substrate repertoire of the CTLH^Muskelin^ C-degron pathway led us to uncover a critical role for the hitherto uncharacterized protein AAMP in ribosome biogenesis. The synthesis of ribosomal proteins is challenging owing to their high abundance and physicochemical properties which render them prone to aggregation, necessitating the evolution of a network of dedicated chaperones^60,65,66^. In yeast Sqt1 is thought to bind nascent uL16 co-translationally, shielding its positively charged rRNA-binding residues prior to incorporation into pre-60S subunits^60^. Structural modeling suggests that uL16, ISG20L2 and AEN and are likely to engage AAMP in an analogous fashion to the uL16-Sqt1 interaction, with a basic α-helix located at or near their N-termini engaging an acidic pocket formed by the WD40 repeats of AAMP. This raises the possibility that these proteins may have to compete for a common binding site on AAMP. As we observed reduced abundance of both uL16 and ISG20L2 upon AAMP depletion, an intriguing possibility is that AAMP could serve to coordinate the rate of rRNA maturation via ISG20L2 with the rate of large subunit maturation via uL16. Thus, we speculate that AAMP may be an important regulatory node controlling both early and late steps of ribosome biogenesis, which may explain why its levels are subject to regulation by the CTLH complex.

## ACKNOWLEDGEMENTS

We are grateful to Gabriela Grondys-Kotarba and Reiner Schulte at the Cambridge Institute for Medical Research Flow Cytometry Core Facility and to the NIHR Cambridge BRC Cell Phenotyping Hub. This work was supported by an Academy of Medical Sciences Springboard Grant (SBF007\100019), an Isaac Newton Trust/Wellcome ISSF/University of Cambridge Joint Research Grant and an ERC Starting Grant (ERC-2024-STG 101160971) to R.T.T. R.T.T. is a Pemberton-Trinity Fellow. I.A.T. is supported by Wellcome (092096) and Cancer Research UK (C6946/A14492); M.P.W. is supported by a Wellcome Discovery Award (309425/Z/24/Z); S.J.E. is supported by the National Institute of Health (AG11085) and is an HHMI investigator; A.J.W. is supported by Cancer Research UK (DRCNPG-Jun24/100002), the Medical Research Council (UKRI1443) and the European Cooperation in Science and Technology (COST) Action Translacore CA21154.

## AUTHOR CONTRIBUTIONS

D.W.G. and R.T.T. conceived the study. D.W.G., S.T., D.E.R., M.Z.L. and Y.D. performed the experiments and analyzed the data, supervised by M.P.W, S.J.E., A.J.W. and R.T.T.. I.A.T. provided essential reagents. D.W.G. and R.T.T. wrote the manuscript.

## DATA AVAILABILITY

Additional data and/or reagents that support the findings of this study are available from the corresponding author upon reasonable request.

## MATERIALS & METHODS

### Cell culture

HEK-293T cells (ATCC CRL-3216) were cultured at 37°C with 5% CO2 in Dulbecco’s Modified Eagle Medium (DMEM) (Merck, #D6429) supplemented with 10% fetal bovine serum (Gibco, #A5256701) plus penicillin and streptomycin (Gibco, #15140122). For total amino acid depletion, the media was replaced with Earle’s Balanced Salt Solution (EBSS) (Gibco, #24010043) supplemented with dialysed fetal bovine serum (Gibco, #A3382001) and vitamins (Gibco, #11120052).

### Chemicals

The following inhibitors were used in this study: Bortezomib (Cambridge Bioscience, #CAY10008822), TAK-243 (Cambridge Bioscience, #HY-100487), MLN4924 (Cambridge Bioscience, #CAY15217), Torin-1 (Stratech Scientific, #A8312-APE), Bafilomycin A1 (ThermoFisher Scientific, #J61835.MCR), Erastin (Cambridge Bioscience, #CAY17754), Tunicamycin (Cambridge Bioscience, #T104) and Buthionine sulfoximine (BSO) (Merck, #19176). All compounds were typically applied to cells for 16 h at a final concentration of 1 μM, apart from Torin-1 which was used at a final concentration of 250 nM. Recombinant human TNF-α was purchased from ThermoFisher Scientific (#PHC3015) and used at a final concentration of 10 ng/ml; recombinant human interferon-ɣ was purchased from Cambridge Bioscience (#229-20150) and used at a final concentration of 50 ng/ml.

### Antibodies

Primary antibodies used were: rabbit α-MAEA (Proteintech, #28363-1-AP), rabbit α-TWA1 (Novus Biologicals, #NBP1-32596), mouse α-Muskelin (Santa Cruz, #sc-398956), mouse α-RanBP9 (Santa Cruz, #sc-271727), rabbit α-RanBP10 (Proteintech, #21107-1-AP), mouse α-ARMC8 (Santa Cruz, #sc-365307), rabbit α-WDR26 (Bethyl, #A302-244A), rabbit α-YPEL5 (Proteintech, #11730-1-AP), rabbit α-AAMP (Abcam, #EPR12369), rabbit α-ISG20L2 (Proteintech, #24639-1-AP), rabbit α-HMGCS1 (Proteintech, #17643-1-AP), mouse α-UBE2H (Santa Cruz, #sc-100620), rabbit α-uL16/RPL10 (Proteintech, #17013-1-AP), rabbit α-uL14/RPL23 (Abcam, #ab264369), mouse α-eS6/RPS6 (Cell Signaling Technology, #2317), mouse α-β-actin (Merck, #A2228), mouse α-vinculin (Merck, #V9131), rabbit α-HA-tag (Cell Signaling Technology, #C29F4), mouse α-FLAG (Merck #F1804), rabbit α-GFP (Abcam, #ab290), mouse α-GFP (Roche, #11814460001) and rabbit α-Phospho-p70 S6 Kinase (Thr389) (Cell Signaling Technology, #9205). HRP-conjugated donkey anti-mouse IgG (#715-035-150) and donkey anti-rabbit IgG (#711-035-152) secondary antibodies were obtained from Jackson ImmunoResearch.

### Plasmids

For individual CRISPR-mediated gene disruption experiments, top and bottom strand oligonucleotides were synthesized (IDT), phosphorylated using T4 PNK (NEB #M0201), annealed by heating to 95°C followed by slow cooling to room temperature, and then ligated into lentiCRISPRv2 (Addgene #52961, kindly deposited by Feng Zhang^67^) cut with BsmBI (NEB #R0739S) using T4 ligase (NEB #M0202). For shRNA-mediated knockdown, hairpins were cloned into the pHR-SIREN-PU6-shRNA-WPRE-PPGK-Puro lentiviral vector (a gift from Prof. Paul Lehner) in an analogous manner utilizing the BamHI and EcoRI restriction enzyme sites. Targeting sequences are detailed in **Table S6**. For individual stability profiling experiments, ORFs were cloned into the pHAGE-GPS vector downstream of GFP using the BstBI and XhoI restriction sites. Exogenous expression of Muskelin mutants was achieved using the pHAGE-PCMV-FLAG expression vector, also via BstBI and XhoI restriction sites. Expression of all other constructs was achieved using pKLV2-based lentiviral vectors, with the pKLV2 sgRNA expression vector (Addgene #67974, kindly deposited by Kosuke Yusa^68^) modified by replacing the U6-sgRNA cassette with the SFFV promoter driving the gene of interest using the MluI and BamHI restriction sites.

### Lentivirus production

Lentiviral packaging was achieved by transfecting HEK-293T cells with the specific lentiviral transfer vector plus a mix of packaging vectors encoding Gag-Pol, Rev, Tat and VSV-G. HEK-293T cells at a confluency of ∼80% were transfected using PolyJet In Vitro DNA Transfection Reagent (SignaGen Laboratories, #SL100688) as per the manufacturer’s protocol. The media was replaced 24 h post transfection and the viral supernatant harvested a further 24 h later. Viral supernatants were centrifuged at 800 x *g* to pellet cell debris and then either directly applied to target cells or stored in single-use aliquots at −80°C.

### Flow cytometry and FACS

Flow cytometry analysis was conducted on an LSR-II instrument using FACSDiva software (BD), with a minimum of 10,000 cells live collected per sample. Subsequent data analysis was performed using FlowJo software (BD). Cell sorting was conducted using an Influx instrument (BD).

### GPS-ORFeome screens

The manufacture of the barcoded ORFeome V8.1 expression library in the pHAGE lentiviral vector has been described previously^69^. The GPS-ORFeome V8.1 library was created by replacing the CMV promoter with a PCMV-DsRed-IRES-GFP cassette using the PI-SceI and I-PpoI restriction enzymes. The resulting library was first packaged into lentiviral particles, and then HEK-293T cells were transduced at a multiplicity of infection (MOI) of ∼0.2 (20% DsRed^+^ cells) at sufficient scale to ensure that at least 10 million cells were transduced. After 48 h, untransduced cells were eliminated through puromycin selection for 3 days. At 7 days post-transduction, stability profiling was performed by partitioning the population into six equal bins based on the GFP/DsRed ratio. Genomic DNA was extracted from all of the sorted cells using the Gentra Puregene Cell Kit (Qiagen, #158767) and the barcodes associated with each ORF were amplified by 24 cycles of PCR (Q5 Hot Start High-Fidelity DNA Polymerase, NEB #M0493S) using primers **(Table S6)** that anneal to common regions flanking the barcode. A pool of eight “staggered” forward primers, each differing in length by one nucleotide, were used in introduce nucleotide diversity for the Illumina platform. Products were purified using a spin column (Qiagen QIAquick PCR Purification Kit, **#**28104) and then 200 ng used as the template for seven additional cycles of PCR designed to add Illumina P5 and P7 adaptors and indexes to allow sample multiplexing. Samples were again purified by using a spin column, quantified by Nanodrop spectrophotometry, and mixed evenly. The final pool was purified by gel extraction from a 2% agarose gel (Qiagen QIAEX II Gel Extraction Kit, #20021) and sequenced using an Illumina NovaSeq 6000 instrument.

### CRISPR/Cas9 genetic screen

The Sabatini/Lander genome-wide CRISPR sgRNA library^70^ was obtained from Addgene (#100000010) and subcloned into the pKLV2 sgRNA expression vector (Addgene #67974, kindly deposited by Kosuke Yusa^68^). The library was packaged into lentiviral particles and HEK-293T cells stably expressing GPS-ZMYND19 and Cas9 transduced at a multiplicity of infection of ∼0.2 at sufficient scale to obtain at least 100-fold representation of the sgRNA library. Puromycin selection was used to eliminate untransduced cells, commencing 48 h post-transduction. On day 7 post-transduction, the top 1% of cells exhibiting stabilization of GPS-ZMYND19 (based on the GFP/DsRed ratio) were isolated by FACS. The sorted cells were returned to culture, and the process repeated 5 days later to further purify the population; genomic DNA was then extracted from both the sorted population and from an unsorted reference population cultured for the same amount of time. Illumina libraries were prepared using an analogous procedure to that described above; primer sequences are detailed in **Table S6**.

### Saturation mutagenesis stability profiling of ZMYND19

For each of the C-terminal 36 residues of ZMYND19 (residues 192-227), an oligonucleotide pool was designed such that each residue was replaced with all other possible amino acids. To maximize sequence diversity between library members, a weighted random selection process was used to assign all codons based on codon usage frequency across the human proteome. The oligonucleotide pool was synthesized by Twist Bioscience. The mutagenized region was amplified by PCR using primers binding to common flanking regions present on each oligonucleotide, purified via extraction from an agarose gel (QIAEX II gel extraction kit, QIAGEN #20021), and then inserted into the GPS lentiviral vector cut BstBI and XhoI along with a second PCR fragment encoding the first 191 residues of wild-type ZMYND19 using the Gibson assembly method (NEBuilder HiFi DNA Assembly Master Mix, NEB #E2621S). The product of the assembly reaction was then concentrated using SPRIselect beads (Beckman Coulter, #B23317) and 2 μl electroporated into ElectroMAX DH10B *E. coli* (ThermoFisher Scientific, #18290015). Bacteria were recovered in S.O.C. Medium (ThermoFisher Scientific, #15544034), incubated for 1 h at 37°C with shaking, and then grown overnight on LB agar plates supplemented with ampicillin (100 μg/ml) at 30°C overnight. Bacterial lawns were scraped from the plates and plasmid DNA extracted using the GenElute HP Plasmid Midiprep Kit (Merck, #NA0200). Following packaging into lentiviral particles, the library was introduced into HEK-293T cells at a multiplicity of infection of ∼0.2 at sufficient scale to achieve approximately 1000-fold representation of the library. Following puromycin selection, the resulting cells were partitioned into three stability bins based on the GFP/DsRed ratio. Genomic DNA was extracted each sorted population using the Quick-DNA Microprep Kit (Zymo Research, #D3021), and the mutant constructs in each stability bin quantified by two rounds of PCR amplification and Illumina sequencing as described above.

### Bioinformatic analysis of genetic screen data

To generate raw count tables quantifying the abundance of each construct in each sample, raw fastq sequence reads were first trimmed of constant flanking sequences using Cutadapt (version 4.1) and then aligned to a reference index detailing all members of the library using Bowtie 2 (version 2.4.5). For the combined GPS-ORFeome screens, a protein stability index (PSI) metric was generated reflecting the distribution of sequencing reads across the six stability bins, given by the sum of multiplying the proportion of reads in each bin by the bin number, resulting in a stability score between 1 (maximally unstable) and 6 (maximally unstable). For the genome-wide CRISPR/Cas9 genetic screen, the raw count table was analyzed using the MAGeCK algorithm^71^ to identify genes targeted by multiple sgRNAs enriched in the sorted population. For the site-saturation stability profiling of the ZMYND19 C-terminus, a PSI score between 1 (maximally unstable) and 3 (maximally stable) was generated for each mutant, reflecting the distribution of sequencing reads across the three stability bins.

### Structure prediction

Structural models were generated using AlphaFold 3 run via the AlphaFold Server (https://alphafoldserver.com/) and further analyzed using UCSF ChimeraX^72^.

### TMT whole cell proteomics

Approximately 5 million HEK-293T cells were harvested in ice-cold PBS and washed once with ice-cold PBS. Lysis buffer composed of 6 M guanidine HCl (ThermoFisher Scientific, #24115) and 50 mM HEPES (Merck, #H0887) was then added for 10 min at room temperature, before the lysates were sonicated. Cell debris was removed by two sequential centrifugation steps (21000 x *g*, 10 min, 4°C). The samples were then incubated at room temperature for 20 min following the addition of dithiothreitol (DTT) at a final concentration of 5 mM. Cysteine residues were then alkylated through the addition of 15 mM iodoacetamide and incubation for 20 min at room temperature in the dark. Excess iodoacetamide was quenched with DTT for 15 min. Samples were diluted with 200 mM HEPES pH 8.5 to reach a guanidine concentration of 1.5 M, followed by digestion with LysC protease (1:100 protease-to-protein ratio) at room temperature for 3 h. Samples were then further diluted with 200 mM HEPES pH 8.5 to reach a guanidine concentration of 0.5 M, before trypsin was added (1:100 protease-to-protein ratio) overnight at 37°C. The reaction was quenched with 5% formic acid and the samples centrifuged (21000 x *g*, 10 min, 4°C) to remove undigested protein. Peptides were subjected to C18 solid-phase extraction (Sep-Pak, Waters) and vacuum-centrifuged to near-dryness. TMT labeling, MS data acquisition and data analysis were performed exactly as described previously^73^, except that no offline HpRP fractionation step was included. The TMT labelling strategy was as follows: sgControl1_rep1, 126; sgControl1_rep2, 131N; sgControl2_rep1, 130N; sgControl2_rep2, 134C; sgTWA1_rep1, 129C; sgTWA1_rep2, 134N; sgMuskelin_rep1, 127N; sgMuskelin_rep2, 131C.

### Immunoblotting

Cells were washed once in phosphate-buffered saline (PBS) and then lysed in 1% sodium dodecyl sulfate (SDS) buffer supplemented with 1:200 Benzonase (Merck, #E1014) for 20 minutes at room temperature. Following addition of Laemmli Buffer (Bio-Rad, #1610747), lysates were heated at 70°C for 10 minutes. Proteins were then separated via polyacrylamide gel electrophoresis using 4-12% Bis-Tris gels (Merck, #MP41G12). Proteins were transferred using the Trans-Blot SD Semi-Dry Transfer System (Bio-Rad) onto PVDF membrane (Merck, #IPVH00010) activated in methanol. After blocking for at least 30 min in 5% skimmed milk powder (Merck, #70166) dissolved in PBS plus 0.1% Tween-20 (PBS-T), membranes were probed with primary antibody at 4°C overnight. Following three 5-minute washes in PBS-T, HRP-conjugated secondary antibodies were applied for 40 minutes at room temperature. After a further five 5-minute washes, reactive bands were visualized using SuperSignal West Detection Reagents (ThermoFisher Scientific, #32106, #34580 and #34076) and images collected from ChemiDoc Imaging System (BioRad). Images were processed using GNU Image Manipulation Platform (GIMP).

### Immunoprecipitation

Approximately 10 million cells were harvested and washed twice with PBS. Cells were lysed using 1% IGEPAL (Merck, #I8896) supplemented with 1:200 Benzonase (Merck, #E1014) plus a cOmplete protease inhibitor tablet (Sigma, #11873580001) on ice for 30 min with periodic inversion. Insoluble material was pelleted by centrifugation (21000 x *g,* 10 min, 4°C) before the supernatant was incubated for 2 h on a rotating wheel with 1 μg antibody plus 10 μl of either Protein A (ThermoFisher Scientific, #10001D) or Protein G magnetic beads (ThermoFisher Scientific, #10003D), or, in the case of the immunoprecipitation of GFP-fusion proteins, 5 μl ChromTek GFP-Trap magnetic beads (Proteintech, #gtma). In each case, the beads were then washed three times in lysis buffer, with 5 min intervals on a rotating wheel between each wash. Bound proteins were eluted by the addition of 1X Laemmli buffer (Bio-Rad, #1610747) supplemented with β-mercaptoethanol followed by heating to 70°C for 10 min. Samples were then further processed by immunoblot as described above.

### Crystal violet staining

Cells were harvested into 5 ml FACS tubes and washed twice with 3 ml PBS. Cells were fixed by adding 250 μl 100% ethanol, incubated at 4°C for 20 min, and then washed twice with 3 ml PBS. Staining was achieved by resuspending the cell pellet in 100 μl 1X crystal violet solution (Pro-Lab Diagnostics, #PL.8000) followed by incubation at room temperature for 30 min. Cells were then washed three times with 3 ml PBS before 150 μl 100% ethanol was added to solubilize bound dye. Five minutes later, the resulting solutions were transferred into 96-well plates and absorbance at 595 nm measured on an Omega FLUOstar (BMG Labtech) instrument.

### Imaging

Cells were imaged using a Zeiss Primovert Inverted Phase Contrast Microscope Ph1/0.3 at 10x magnification via a NexYZ 3-axis Universal Smartphone Adapter (Celestron).

### Polysome profiling

Ribosomal subunits were separated by sucrose density gradients essentially as described previously^74–76^. Approximately 30 million cells were treated with 100 μg/ml cycloheximide (Sigma) for 15 min at 37°C, washed twice with PBS supplemented with 100 μg/ml cycloheximide, and then lysed in lysis buffer composed of 20 mM HEPES at pH 7.4, 50 mM KCl, 5 mM magnesium acetate, 0.5% [v/v] IGEPAL® CA-630 (Sigma), 0.5% [w/v] deoxycholate (Sigma), with an EDTA-free protease inhibitor tablet (Roche) plus 100 μg/ml cycloheximide (Sigma) and 500 U/mL RNase inhibitor (RNaseOUTTM, Invitrogen). Following a 15 min incubation on ice, lysates were clarified by centrifugation (20000 x *g*, 10 min, 4°C). Sucrose gradients were prepared using a Biocomp Gradient Master according to the manufacturer’s manual. Equal amounts (typically 3.0 A260 nm unit) of sample were loaded onto 10% – 40% (w/v) sucrose gradients in 14 ml gradient buffer (20 mM HEPES pH 7.4, 50 mM KCl, 5 mM magnesium acetate with complete EDTA-free protease inhibitors (Roche) plus 100 µg/ml cycloheximide) and centrifuged (284600 x *g*, 2.5 h, 4°C using a Beckmann SW40Ti rotor). Samples were unloaded using a Brandel gradient fractionator. Polysome profiles were detected using an ÄKTAprime Plus system (GE Healthcare) and 0.5 ml fractions collected. Proteins were subsequently precipitated with 20% (w/v) trichloroacetic acid for immunoblot analysis.

## SUPPLEMENTARY FIGURE LEGENDS

**Supplementary Figure 1.**
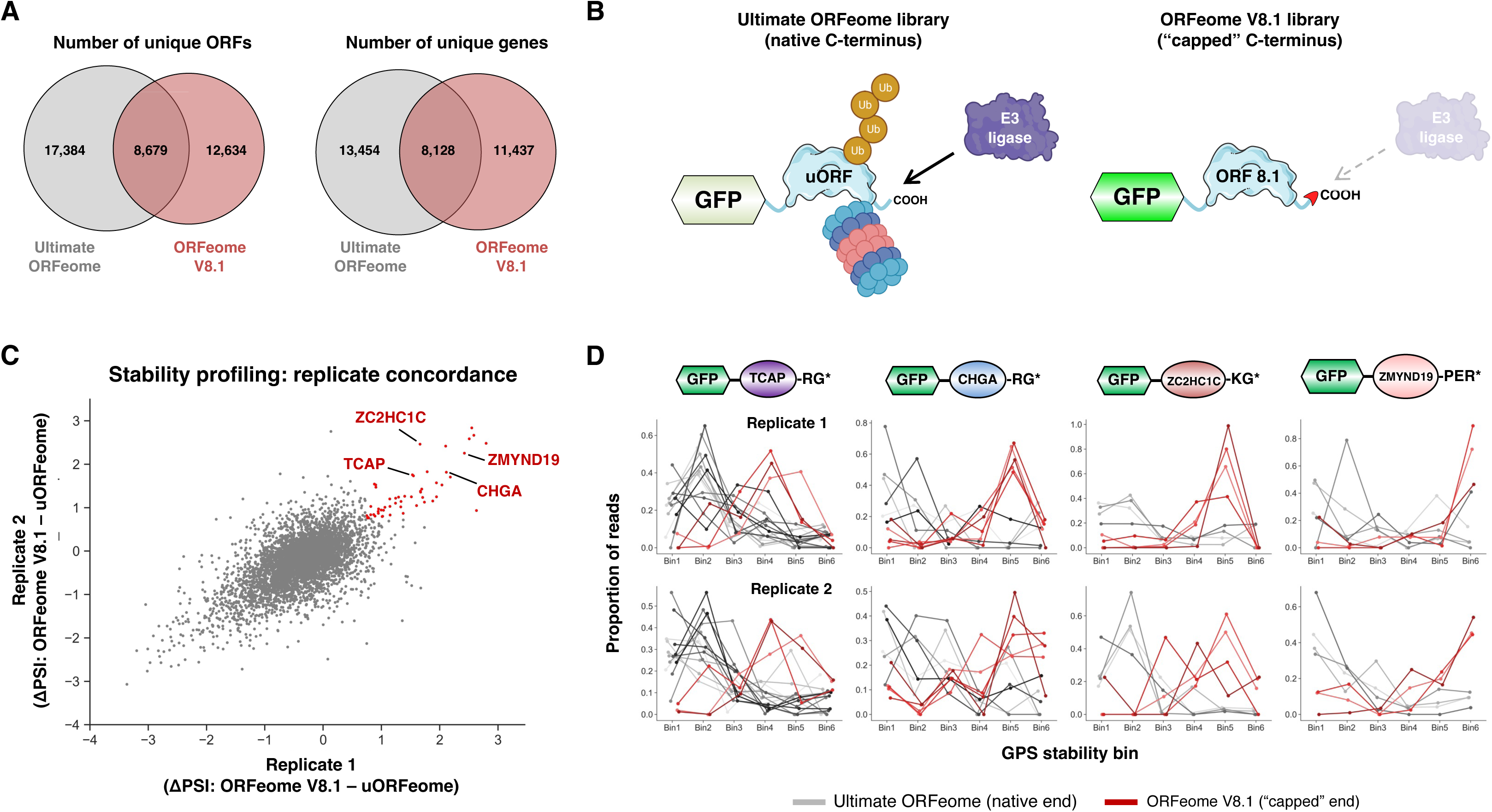
Comparative stability profiling identifies full-length human proteins bearing C-terminal degrons. **(A)** Venn diagrams representing the overlap between the constituent members of the two ORFeome libraries. **(B)** Schematic depicting the rationale underpinning the comparative screening approach. Should an E3 ligase target a substrate for ubiquitination and degradation via a degron located at its extreme C-terminus (left), we reasoned that “capping” of the C-terminus via the addition of an invariant 9 amino acids in the ORFeome V8.1 expression vector (right) would prevent E3 ligase recognition and result in stabilization of the substrate. **(C)** Assessing the concordance between replicate screens. Each dot depicts the performance (ΔPSI, the difference in stability when assayed as part of the ORFeome V8.1 library compared to the Ultimate ORFeome library) of each protein detected in both libraries across duplicate experiments. Red dots indicate the 52 proteins exhibiting concordant stabilization >0.75 PSI units as part of the ORFeome V8.1 library. **(D)** Assessing the screen performance of individually barcoded replicates for the selected substrates. Each ORF in the libraries is represented by an average of five independent barcodes, thus providing internal replicates; screen profiles (depicting the distribution of sequencing reads across the six stability bins) for each individually barcoded replicate are shown, when assayed as part of the Ultimate ORFeome (gray traces) compared to the ORFeome V8.1 (red traces).

**Supplementary Figure 2.**
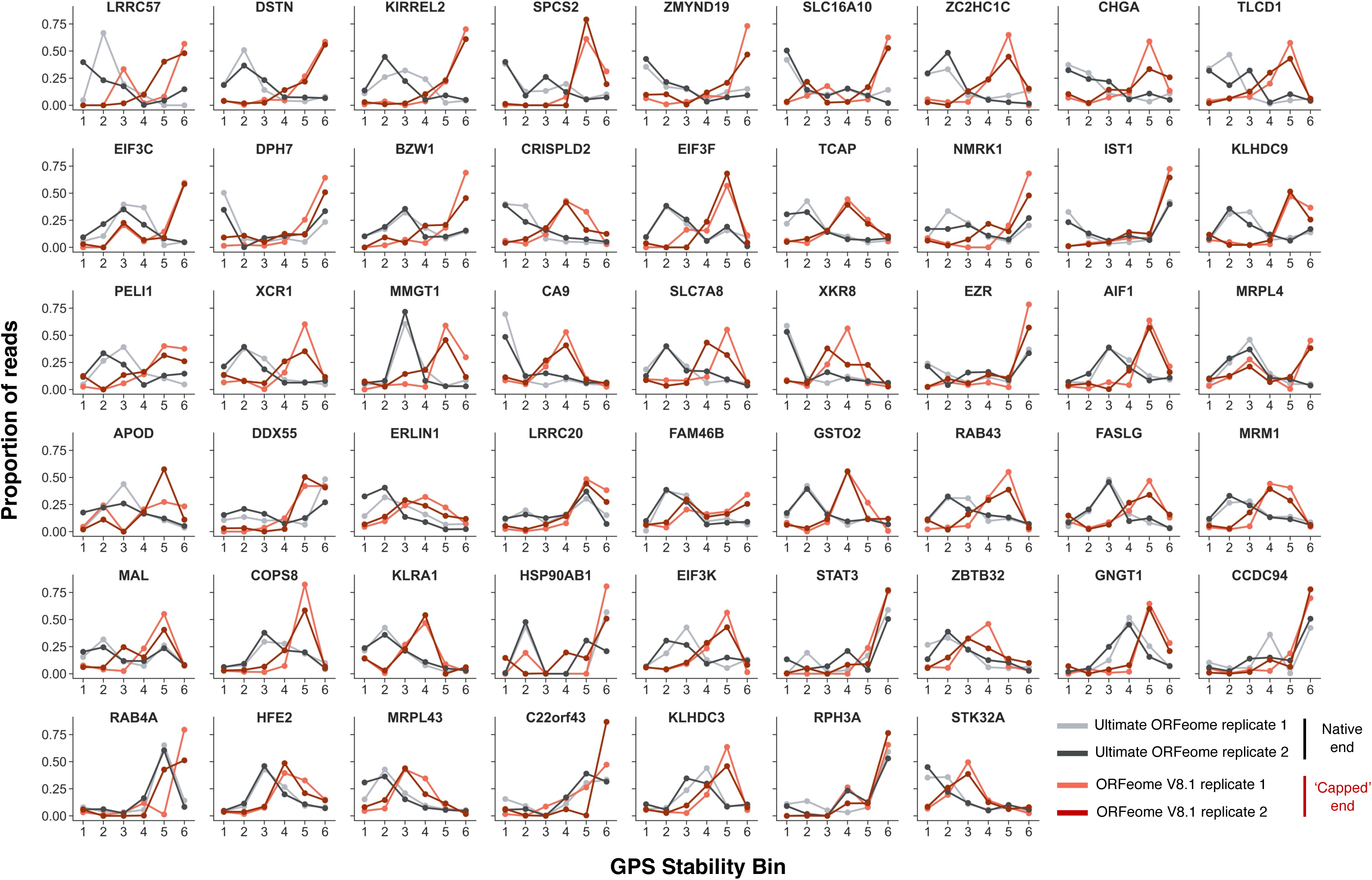
Candidate proteins harboring C-terminal degrons. Screen profiles are shown for all 52 proteins exhibiting concordant stabilization when assayed as part of the ORFeome V8.1 library as compared to the Ultimate ORFeome library across duplicate screens.

**Supplementary Figure 3.**
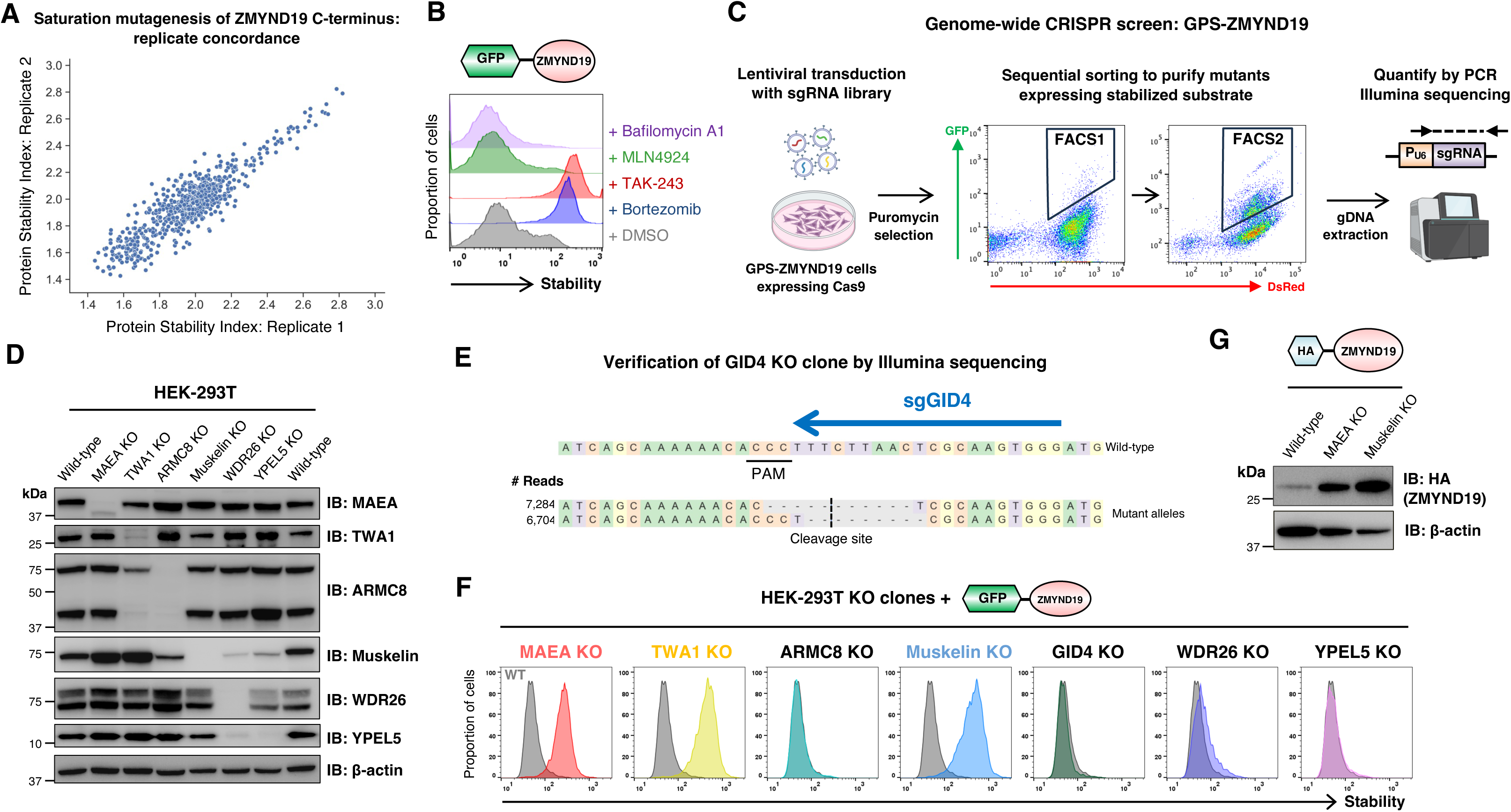
Defining the machinery required for the degradation of ZMYND19. **(A)** Saturation mutagenesis of the ZMYND19 C-terminus yields highly concordance stability scores. The scatterplot reflects the protein stability index metric (1 = maximally unstable, 3 = maximally stable) for each ZMYND19 mutant across two replicate experiments. **(B)** Ubiquitin-dependent proteasomal degradation of ZMYND19. HEK-293T cells expressing GPS-ZMYND19 were treated with the indicated inhibitors and assessed by flow cytometry 8 hours later. Treatment with the proteasome inhibitor Bortezomib and the E1 inhibitor TAK-243 resulted in ZMYND19 stabilization, whereas the pan-Cullin inhibitor MLN4924 and the lysosomal inhibitor Bafilomycin A1 did not. **(C)** Schematic representation of a genome-wide CRISPR screen designed to identify the machinery required for ZMYND19 degradation. **(D-E)** Validation of knockout clones lacking CTLH subunits generated by CRISPR/Cas9-mediated gene disruption. Validation was performed by immunoblot **(D)** with the exception of the GID4 KO clone, which, owing to the lack of effective commercial antibodies, was assessed by Illumina sequencing **(E)**. **(F)** Assessing the stability of ZMYND19 in the KO clones lacking CTLH subunits. GPS-ZMYND19 was expressed in clones of the indicated genotypes and its stability assayed by flow cytometry. **(G)** An HA-tagged ZMYND19 construct is more abundant in MAEA KO and Muskelin KO clones, as assessed by immunoblot.

**Supplementary Figure 4.**
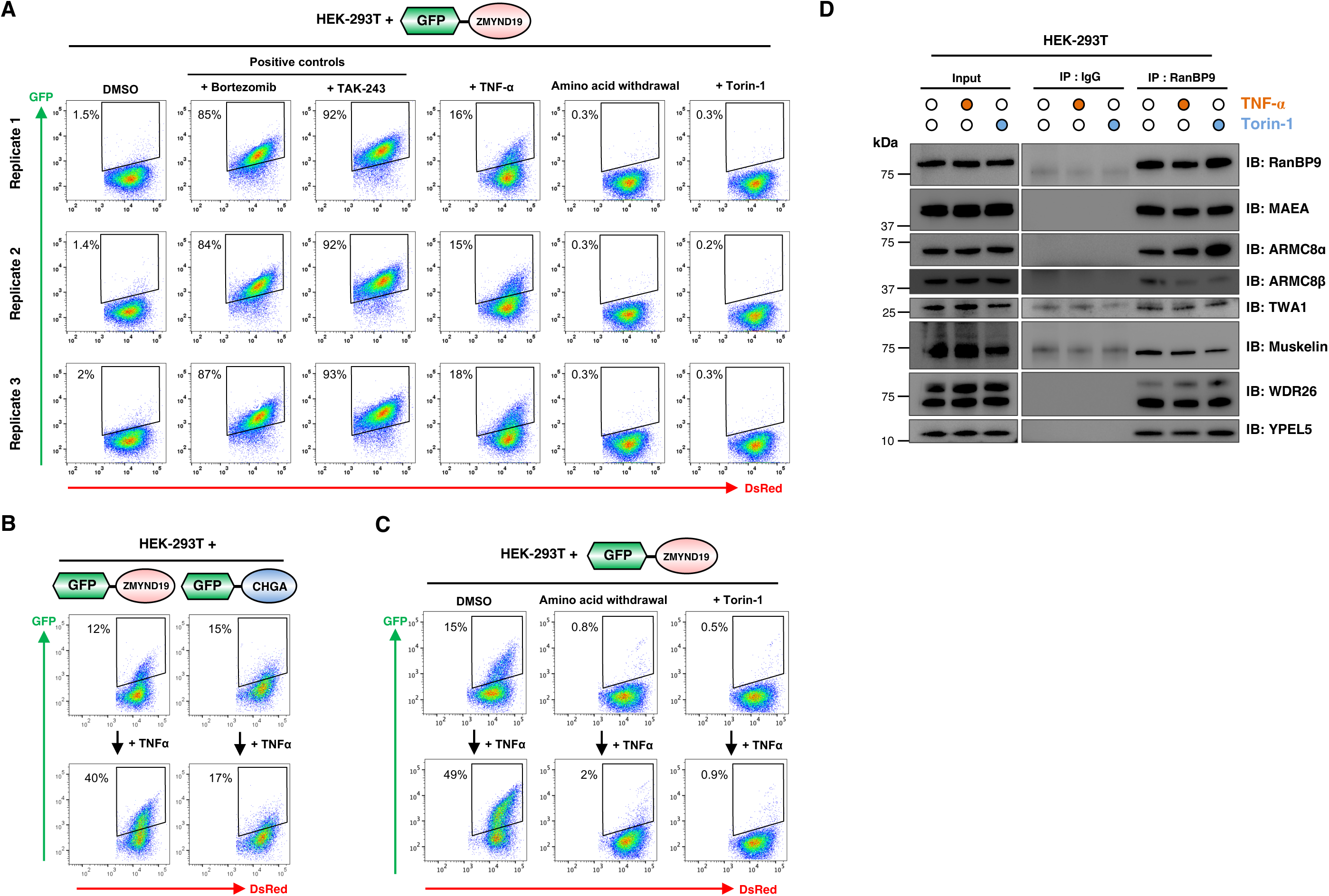
Conditional regulation of ZYMND19 degradation. **(A)** Reproducible effects of TNF-α stimulation and mTOR inhibition on ZMYND19 stability. HEK-293T cells expressing ZMYND19 in the context of the GPS expression vector were subjected to the indicated treatments in triplicate, and the effect on the stability of ZMYND19 assayed by flow cytometry. **(B)** TNF-α stimulation does not broadly affect proteasomal degradation. HEK-293T cells were transduced with GPS vectors encoding ZMYND19 and CHGA, treated with TNF-α overnight and then analyzed by flow cytometry. **(C)** Amino acid withdrawal or mTOR inhibition reproducibly overrides the effects of TNF-α and promotes ZMYND19 degradation. A replicate experiment of the one performed in Fig. 4E is shown. **(D)** Assessing the effects of TNF-α stimulation and mTOR inhibition on CTLH complex assembly. CTLH was immunoprecipitated from HEK-293T cells treated with either TNF-α or Torin-1 using an antibody against RANBP9, and co-immunoprecipitating CTLH subunits assessed by immunoblot.

**Supplementary Figure 5.**
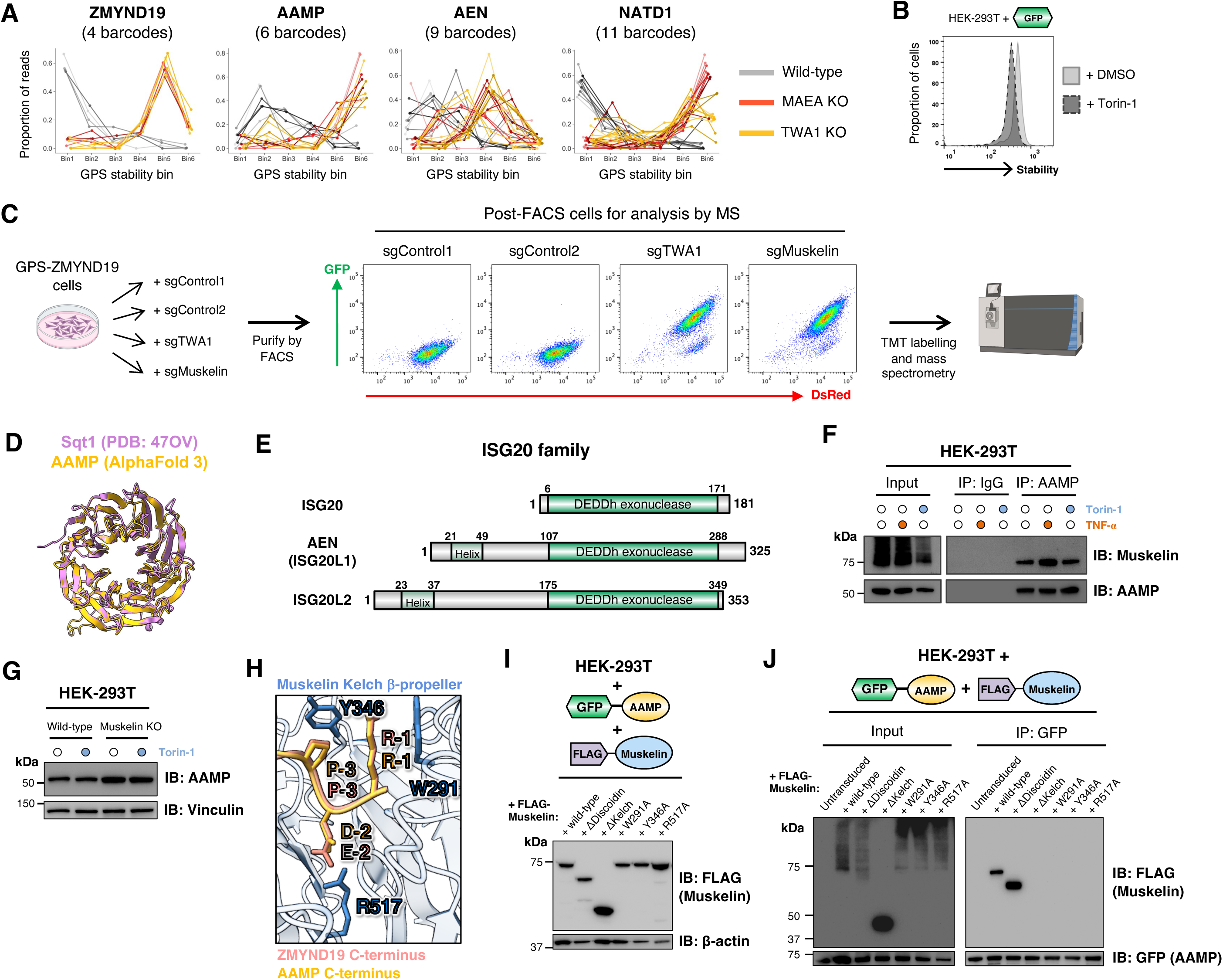
Identifying additional substrates of the CTLH^Muskelin^ C-degron pathway. **(A)** Performance of individually barcoded replicates for the CTLH substrates identified by the stability profiling screen. **(B)** A small reduction in stability is observed following overnight Torin-1 treatment when GFP alone is expressed in the context of the GPS vector. **(C)** Purifying cells lacking CTLH subunits for proteomic analysis. HEK-293T cells expressing GPS-ZMYND19 were transduced in duplicate with Cas9 and sgRNAs targeting the indicated genes; cells lacking a functional CTLH complex stabilize ZMYND19 and can therefore be isolated by sorting for GFP^+^ cells. Two intron-targeting sgRNAs were used in parallel as negative controls, which were also sorted in an equivalent manner (targeting DsRed^+^/GFP^-^cells). **(D)** Overlay of the crystal structure of Sqt1 (pink) and the AlphaFold 3-predicted structure of AAMP (gold). **(E)** Schematic representation of the ISG20 family of proteins. **(F)** The interaction between Muskelin and AAMP is preserved following treatment with Torin-1 or TNF-α. **(G)** The effect of Torin-1 treatment and Muskelin knockout on the abundance of endogenous AAMP, as assessed by immunoblot. **(H-J)** The C-terminus of AAMP is recognized by the β-propellor formed by the kelch repeats of Muskelin. **(H)** Superimposition of AlphaFold 3 structure predictions suggests that Muskelin engages the C-termini of ZMYND19 and AAMP in an analogous manner. **(I)** Expression levels of the indicated FLAG-tagged Muskelin mutants as assessed by immunoblot. **(J)** Muskelin mutants affecting the predicted binding interface abolish the association between AAMP and Muskelin. The indicated FLAG-tagged Muskelin mutants were expressed in cells expressing GPS-AAMP, and binding assessed by immunoprecipitation with an anti-GFP nanobody followed by FLAG immunoblot.

**Supplementary Figure 6.**
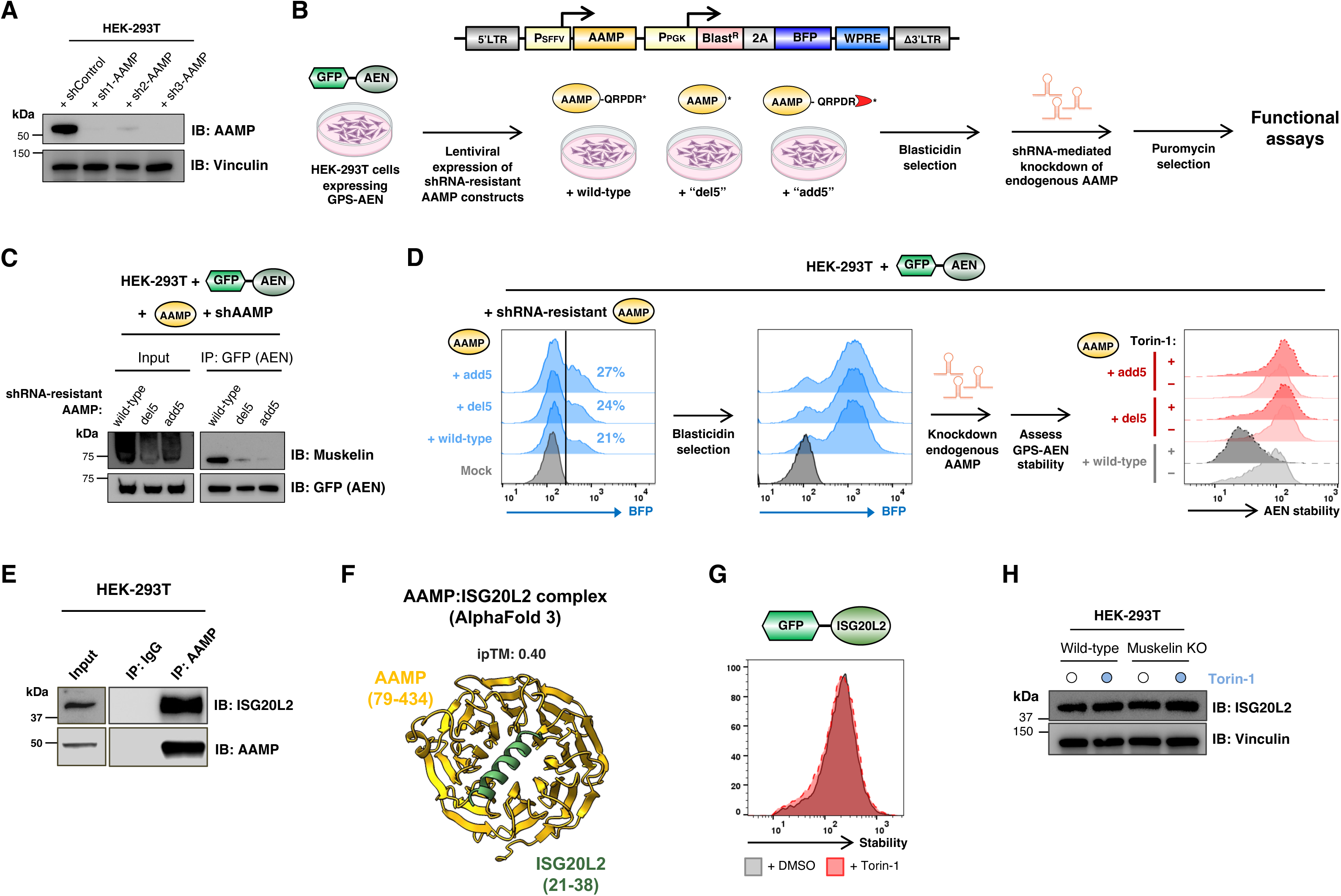
Collateral degradation of the AAMP interacting partner AEN by the CTLH^Muskelin^ C-degron pathway. **(A)** Validation of efficient shRNA-mediated depletion of AAMP. HEK-293T cells were transduced with lentiviral shRNA expression vectors and the extent of AAMP depletion four days post-transduction was measured by immunoblot. The two most efficient shRNAs, sh1-AAMP and sh3-AAMP, were used for subsequent experiments. **(B-D)** AEN is subject to collateral degradation by CTLH^Muskelin^ via AAMP. **(B)** Schematic representation of the experimental setup to determine the impact of manipulating the C-terminus of AAMP on the CTLH-mediated degradation of AEN. Owing to the essentiality of AAMP (see Fig. 7), we performed the genetic complementation assay by first introducing shRNA-resistant AAMP constructs into cells expressing GPS-AEN, followed by shRNA-mediated knockdown of endogenous AAMP. **(C)** The association between AEN and Muskelin requires the C-terminus of AAMP. Immunoprecipitation of GFP-tagged AEN pulled-down Muskelin when wild-type AAMP was present, but to a lesser extent in the presence of mutant versions of AAMP bearing either a truncated (“del5”) or capped (“add5”) C-terminus. **(D)** The degradation of AEN stimulated by mTOR inhibition requires the C-terminus of AAMP. The indicated AAMP constructs were introduced into cells expressing GPS-AEN at single copy, as assessed by flow cytometry measurement of BFP^+^ cells (left panel). Following blasticidin selection (center panel), the cells were transduced with shRNA expression vectors to deplete endogenous AAMP, and, following overnight treatment with Torin-1, the stability of AEN was then assayed by flow cytometry (right panel). **(E-F)** AAMP interacts with ISG20L2. **(E)** Co-immunoprecipitation of ISGL20L2 with AAMP, as detected by immunoblot. **(F)** AlphaFold 3 model of the ISG20L2-AAMP interaction, which suggests a basic α-helix located at the N-terminus ISG20L2 may be accommodated by an acidic pocket formed by the WD40 repeats of AAMP. **(G-H)** ISG20L2 is not a substrate of collateral degradation by the CTLH^Muskelin^ C-degron pathway. The stability of ISG20L2 is not affected by Torin-1 treatment when expressed in the context of the GPS expression system **(G)**, and the abundance of endogenous ISG20L2 is not affected upon ablation of Muskelin **(H)**.

**Supplementary Figure 7.**
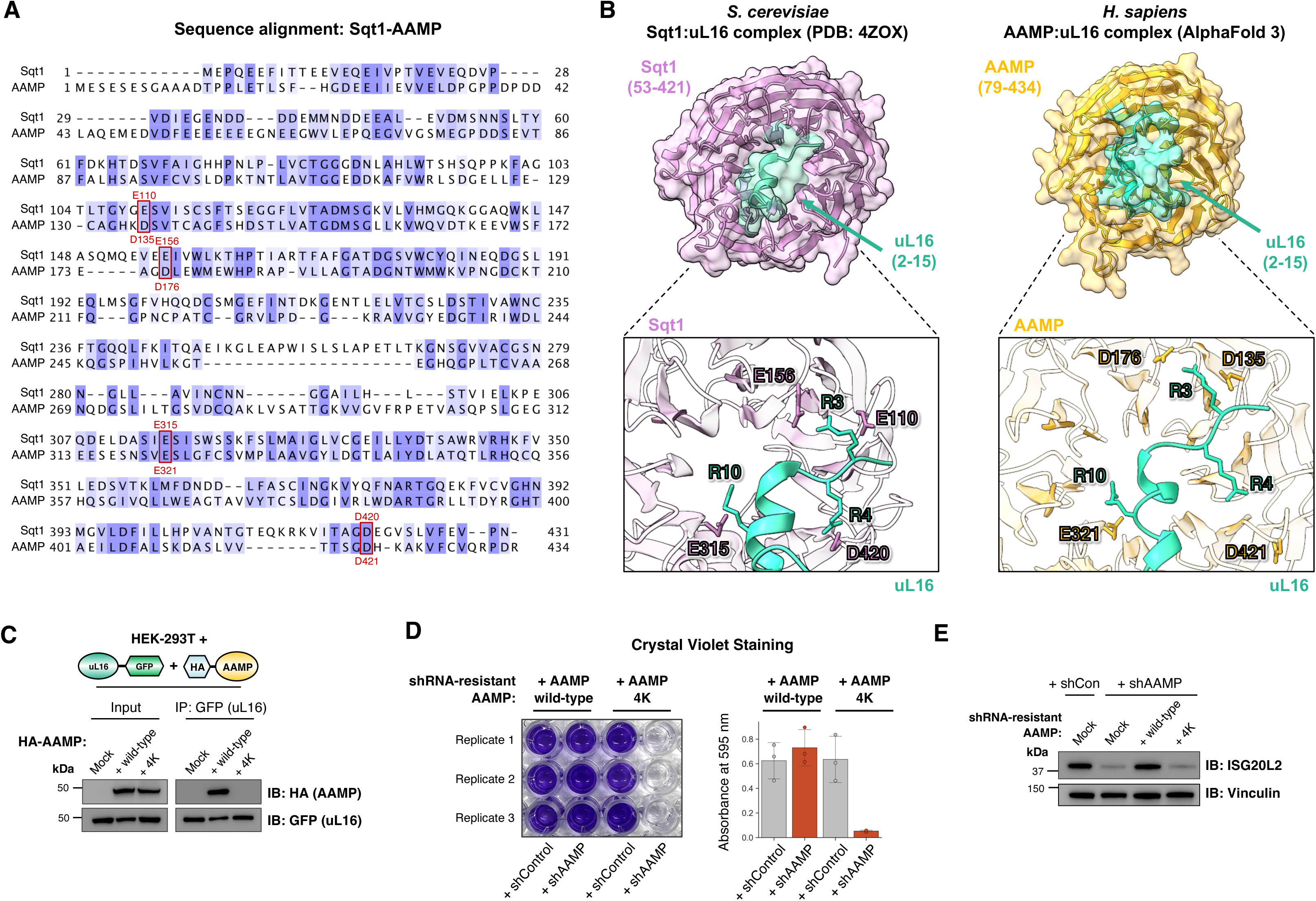
AAMP binds uL16 and ISG20L2 to regulate ribosome biogenesis. **(A)** Sequence alignment of *S. cerevisiae* Sqt1 and *H. sapiens* AAMP. Blue shading reflects the degree of amino similarity. The acidic residues mutated in subsequent experiments are highlighted in red boxes. **(B-D)** Recognition of uL16 by AAMP is necessary for cell viability. **(B)** Design of the AAMP “4K” charge-swap mutant. Analysis of the Sqt1-uL16 interface (left) highlights four acidic residues contacting uL16; the corresponding acidic residues are also predicted to contact uL16 in the AlphaFold 3 model of the AAMP-uL16 interaction (right). **(C)** The 4K charge-swap mutant of AAMP failed to associate with uL16, as assessed by co-immunoprecipitation experiments, and, unlike wild-type AAMP, could not restore cell viability upon knockdown of endogenous AAMP **(D)**. **(E)** Reduced ISG20L2 abundance upon AAMP knockdown. As was the case for uL16 (Fig. 7J), AAMP depletion reduced ISG20L2 protein levels; this effect could be reversed upon exogenous expression of an shRNA-resistant wild-type AAMP, but not by the 4K charge-swap AAMP mutant.

**Supplementary Figure 8.**
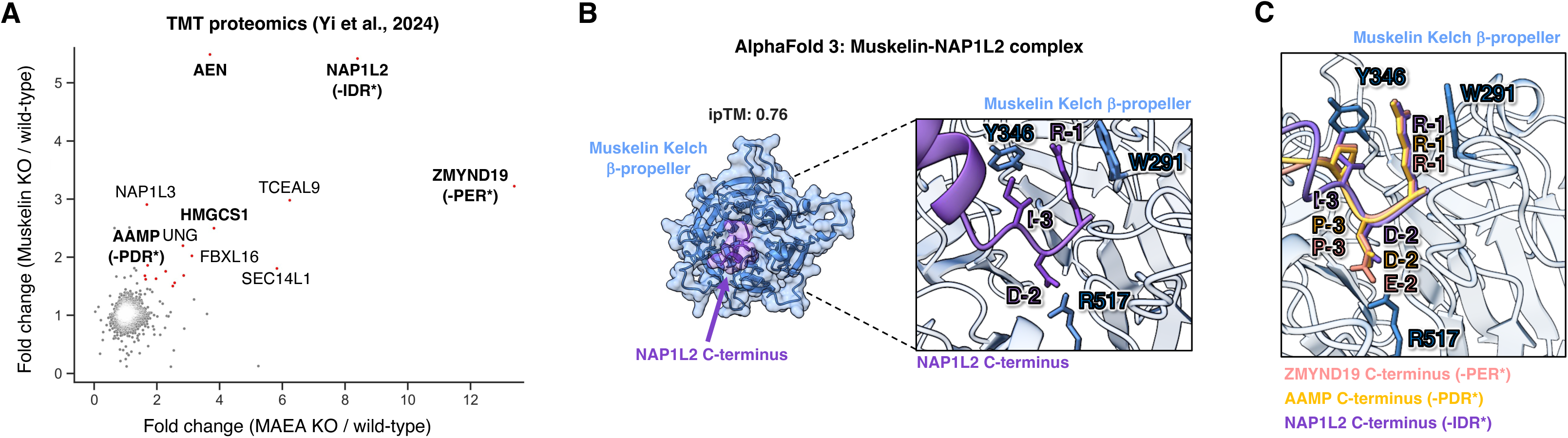
NAP1L2 is a candidate substrate of the CTLH^Muskelin^ C-degron pathway. **(A)** Scatterplot representing TMT proteomics data from. Substrates discussed in this study are shown in bold font. **(B)** AlphaFold 3 prediction of the interaction between the C-terminus of NAP1L2 and the β-propellor formed by the kelch repeats of Muskelin. **(C)** Superimposition of AlphaFold 3 structural predictions, showing that the C-terminus of NAP1L2 is predicted to contact the same residues in the β-propellor formed by the kelch repeats of Muskelin as ZMYND19 and AAMP.

## SUPPLEMENTARY TABLE LEGENDS

**Supplementary Table 1 | Comparative stability profiling screens to identify full-length proteins bearing C-terminal stability determinants.** ORF-level data showing the corrected read counts across the six stability bins for each ORF detected common to both libraries across replicate experiments.

**Supplementary Table 2 | Saturation mutagenesis of the ZMYND19 C-terminus defines a C-terminal degron motif.** Raw data is provided for all ZMYND19 mutants detected in the stability profiling screen. (PSI, protein stability index; 1 = maximally unstable, 3 = maximally stable).

**Supplementary Table 3 | A genome-wide CRISPR screen to identify the machinery required for ZMYND19 degradation.** The output from the MAGeCK algorithm is shown.

**Supplementary Table 4 | Stability profiling identifies additional CTLH substrates.** ORF-level data showing the corrected read counts across the six stability bins for each ORF detected across wild-type (WT), MAEA KO and TWA1 KO cells.

**Supplementary Table 5 | Identifying CTLH^Muskelin^ substrates by TMT mass spectrometry.** Raw data for all proteins quantified is shown.

**Supplementary Table 6 | Oligonucleotide sequences.**

